# Environmental volatility amplifies Pavlovian control in decision-making

**DOI:** 10.64898/2026.09.12.751157

**Authors:** Claudio Danti, Luigi A.E. Degni, Lorenzo Mattioni, Marco Badioli, Francesca Starita, Emilio Cartoni, Sara Garofalo, Giuseppe di Pellegrino

## Abstract

Volatile environments require continuous updating of action values, making instrumental knowledge less informative. Under such conditions, Pavlovian control may offer a more stable alternative, since the value attributed to reward-predictive cues is not altered by continuously updated instrumental predictions. Using a modified Pavlovian-to-Instrumental Transfer paradigm in two preregistered experiments, we test the hypothesis that increased volatility promotes the recruitment of Pavlovian control, whereby reward-predictive cues bias choice to the predicted outcome, simplifying (and possibly overcoming) the high computational cost of the instrumental prediction. Crucially, using entropy-based metrics we demonstrate that Pavlovian control is not unitary, but can instead respectively increase or reduce entropy depending on whether the choice outcome is present or not. Together, these findings show that volatility redirects control in Pavlovian–instrumental arbitration toward the cues, rather than merely adding noise to behaviour.

## Introduction

Adaptive behaviour requires learning which actions lead to desirable outcomes while tracking how reliable those instrumental action–outcome relationships are over time (Rangel et al., 2008). Natural environments are inherently uncertain, and contingencies may shift abruptly, rendering previously learned actions suboptimal. Environmental volatility, i.e., the rate at which action-outcome contingencies unexpectedly change, represents one fundamental form of uncertainty that the brain must estimate and respond to (Soltani & Izquierdo, 2019; Yu & Dayan, 2005).

When volatility is high, computational and behavioural studies consistently show that agents prioritize learning from recent over distal outcomes, as reflected in higher learning rates (Behrens et al., 2007; Blain & Rutledge, 2020; Massi et al., 2018). Even with this adaptation, however, volatile environments impose an unavoidable cost: because contingencies keep changing, recent outcomes provide a noisier guide to which action is currently optimal, reducing choice accuracy and increasing uncertainty about the best course of action (Farashahi et al., 2017; Soltani et al., 2021; Soltani & Koechlin, 2022). This uncertainty is captured by higher entropy of choice, an index of how consistently behaviour is distributed across the available actions (Trepka et al., 2021; Woo, et al., 2025a; Woo et al., 2023).

One way to contain this cost would be to resort to a system whose responses are triggered by predictive cues rather than by tracked action–outcome contingencies. Pavlovian associations provide exactly such a system: because motivational value is acquired through stimulus-outcome associations rather than action-outcome tracking, changes in instrumental contingencies do not necessarily alter the value attributed to the stimulus. This motivational value also influences instrumental choice. A Pavlovian cue biases action selection toward the action associated with its outcome; for example, a cue predicting chocolate biases choice toward the action that led to chocolate. When Pavlovian and instrumental systems promote the same action, this bias supports efficient decisions; instead, when they promote different actions, the resulting conflict can lead to suboptimal choice (Dayan et al., 2006; Degni et al., 2025; Guitart-Masip et al., 2012; Huys et al., 2011; Swart et al., 2017). An exaggerated form of this cue-driven, suboptimal choice may contribute to conditions such as addiction, impulsivity and compulsive disorders (Everitt & Robbins, 2005; Garofalo & di Pellegrino, 2015; Marzuki et al., 2024; Sommer et al., 2017).

To date, how volatility shapes the balance between instrumental and Pavlovian control over action selection has not been directly investigated. A long-standing computational principle holds that the brain arbitrates between competing systems of valuation (Courville et al., 2006; Daw et al., 2005). Under uncertainty, control shifts toward whichever system (Pavlovian vs Instrumental), at a given moment, reduces uncertainty the most (Soltani & Izquierdo, 2019; Yu & Dayan, 2005). What matters is therefore the relative informativeness of the two systems, and environmental volatility acts directly on this balance: by frequently reversing action–outcome contingencies, it erodes the informativeness of instrumental knowledge while leaving the predictive value of Pavlovian cues intact. Behaviour should then shift toward cue-driven choice, a shift that becomes more visible whenever a Pavlovian cue and instrumental learning conflict by pointing to different actions, so that following the cue means departing from what instrumental learning currently favours (Boureau et al., 2015; Dayan et al., 2006; Dorfman & Gershman, 2019).

Here, we tested this hypothesis across two preregistered experiments using a modified Pavlovian-to-Instrumental Transfer paradigm in which environmental volatility was manipulated through the frequency of contingency reversals. In Experiment 1, participants first learned the outcomes of two actions (one of which was more frequently rewarded than the other) during an instrumental learning phase, completing one block under a low-volatility context and one under a high-volatility context, each signalled by a different background colour. After a Pavlovian learning phase, in which they learned the associations between the three cues and their outcomes, they then completed a Transfer phase, choosing between the same two actions while Pavlovian cues were presented alongside each choice, with outcome feedback (i.e., participants saw the outcome of each choice). We predicted that, under high volatility, there would be a shift in the balance of control toward the Pavlovian system: when a cue predicted an outcome associated with a different action than the one instrumental learning favoured, participants would follow the cue more often, reducing choice accuracy. Because the two systems compete for control on this condition, we further predicted an increase in the entropy of choice strategy. Experiment 2 examined whether this shift persists in the absence of ongoing reinforcement, by interleaving segments with and without outcome feedback during the Transfer phase, informing participants that rewards could still occur even when no outcome was shown. Without outcome feedback, the instrumental system loses the signal it needs to update, leaving the cue as the only source of value. We therefore predicted that when Pavlovian and instrumental predictions conflict, choice would follow the cue even more consistently, further decreasing accuracy. For entropy, we had no directional prediction: a decrease would indicate that the cue came to dominate choice entirely, whereas an increase comparable to Experiment 1 would indicate that the two systems continued to compete even without feedback.

## Experiment 1

### Methods

#### Participants

Healthy adult volunteers (age ≥ 18 years) were recruited by word of mouth and advertisement. Participants were naïve to the purposes of the experiment and provided written informed consent prior to participation. The study was conducted in accordance with institutional guidelines and the 1964 Declaration of Helsinki and was approved by the Bioethics Committee of the University of Bologna.

The target sample was determined a priori by a pre-registered power analysis (<u>10.17605/OSF.IO/9YGHR</u>) conducted on MorePower 6.0 (Campbell & Thompson, 2012) for the planned 3 (CS: CS+_1_, CS+_2_, CS−) × 2 (Volatility: Low, High) repeated-measures ANOVA. The analysis assumed α = 0.05 (two-tailed), an effect size of (η_p_²) = 0.07 (estimated from our pilot study, N = 16 (Supplementary Fig. S1)) and desired power = 0.80, yielding a required sample of N = 66. Consistent with the pre-registered stopping rule, interim Bayesian repeated-measures ANOVAs were performed after every 10 participants starting at N = 30. Data collection was to stop early if the Bayes factor (BF₁₀) for the CS × Volatility interaction exceeded 10 (strong evidence for H₁) or fell below 1/10 (strong evidence for H₀). Data collection was terminated at N = 50 when the stopping criterion was met (BF₁₀ =1.061 ×10^4^). The final sample comprised 50 participants (30 females; mean age = 24.48, sd = 3.56 years, mean education = 16.46, sd = 1.87 years). All participants were included in the analysis.

#### Experimental Paradigm

We adopted a modified version of the Pavlovian-to-Instrumental Transfer (PIT) paradigm (Degni et al., 2025, 2026). The task comprised three consecutive phases: an Instrumental learning phase, in which participants learned response–outcome associations under varying volatility conditions; a Pavlovian learning phase, in which participants learned cue–outcome associations; and a Transfer phase, in which Pavlovian influences on instrumental responding were tested under two possible volatility contexts (high and low).

An image of a grey slot-machine, embedded within a coloured background that signalled the current context, was presented during all task phases. During the Instrumental and Transfer phases, two background rooms (light-blue and orange) indicated the low and high volatility contexts; the colours were matched for luminance and transparency and colour–context assignment was counterbalanced across participants. During the Pavlovian phase, the slot-machine was presented in a neutral white background room, with no volatility manipulation. The slot-machine comprised two displays (upper and lower) and a central fixation cross. During the Instrumental and Transfer phases, two levers (left/right) appeared on the side of the slot-machine to indicate the available actions, selected via corresponding keyboard keys (i.e., “z” for the left and “m” for the right lever of the slot-machine). A running numeric counter was visible during these phases, providing participants with online feedback about their performance. The task was programmed and executed in OpenSesame v4.0 (Mathôt et al., 2012).

### Instrumental learning phase

In the Instrumental learning phase, participants learned the associations between two responses (R_1_ and R_2_) and two corresponding rewarding outcomes (O_1_ and O_2_) under varying volatility conditions. On each trial, an empty slot-machine was presented (inter-trial interval, ITI, 500–1500ms), after which two levers appeared on the left and right sides of the slot-machine and remained visible until a response was made. Once participants responded, the selected lever dropped and the outcome image was displayed in the lower screen for 1000ms. Outcomes consisted of two distinct food rewards (O₁ and O₂) individually tailored to equate subjective value and motivation, whereas a negative outcome (O–, displayed as an “X”) signalled a loss. A running numeric counter was visible throughout this phase and updated trial by trial increasing by one whenever a food reward was delivered and decreasing by one whenever the non-rewarding outcome (X) was displayed. Of note, it was programmed never to drop below zero to maintain motivational feedback. The counter was reset to zero at the start of each block. Participants were informed that they would gain one point for each rewarded trial and lose one point for each non-rewarded trial, with the total number of points determining the amount of food reward received at the end of the experiment.

Volatility was manipulated across blocks, alternating between low and high volatility contexts. In both volatility contexts the payoff structure followed a probabilistic 80% versus 20% reward schedule: in low-volatility blocks, one lever produced its associated rewarding outcome (O₁ or O₂) with 80% probability and the loosing outcome (O–) with 20% probability, whereas the other lever produced the opposite contingency. Approximately halfway through the block (at a pseudorandom point between trial 45 and 55), the reward contingencies of the two levers were reversed, such that the previously better lever became the worse one and vice versa. In high volatility blocks, the same 80/20 structure applied but contingencies reversed five times per block at pseudo-random intervals (every 10–25 trials) to reduce predictability (Soltani & Izquierdo, 2019). Crucially, reversals were not signalled and thus, subjects had to adjust their choice solely based on reward feedback to maximize their chance of winning a reward. In both contexts, the reversal schedule was generated once and held fixed across participants, so that all participants experienced the same sequence of reversals within each volatility condition. To control for potential sequence effects, the order of volatility contexts (starting with low or high) was counterbalanced across participants.

A series of 100-trial blocks was administered and repeated until the instrumental learning criterion was met. The learning criterion required correctly reporting the association between the two responses and the two outcomes. Learning was assessed after four initial blocks (400 trials: thus, 200 trials per volatility condition); if the criterion had not been attained, participants completed an additional pair of 100-trial blocks (one low and one high volatility block) and learning was re-assessed after each pair. Thus, participants completed a minimum of four blocks and a maximum of eight blocks. All participants met the learning criterion after the minimum of four blocks; therefore, analyses of the Instrumental learning phase included 200 trials per condition for all participants. Each block had an average duration of approximately 6 minutes. Before the Instrumental learning phase, participants completed a liking rating to ensure comparable motivational value of the selected outcomes and conditioned stimuli (CSs). A 9-point Likert scale (0 = “not at all”, 9 = “very much”) was used to rate each food reward on two dimensions: “How much do you usually enjoy eating it?” (general liking) and “How much would you like to eat it now?” (current wanting). The same scale was then presented for each of the three CSs, with the question “How much do you like this stimulus?”, to verify that the stimuli were matched in baseline preference before learning.

### Pavlovian learning phase

In the Pavlovian learning phase, participants learned the associations between three conditioned stimuli (CS+_1_, CS+_2_, CS−), represented by fractal-like images and the same outcomes used in the preceding Instrumental phase (O₁, O₂ and O–). Two stimuli served as CSs+ (CS+_1_ and CS+_2_): on 80% of their presentations, CS+₁ predicted outcome O₁ and CS₂ predicted outcome O₂, while on the remaining 20% they were followed by the non-rewarding outcome (O–, displayed as an “X”) (Degni et al., 2025; Garofalo et al., 2021). The third stimulus (CS–) was always followed by O– on 100% of trials. The assignment of CSs to specific outcomes was counterbalanced across participants.

Because this phase involved passive observation rather than instrumental responding, participants were instructed that the slot machine was “broken” and that no levers would be available. To ensure that the O– outcome was not perceived as a loss, participants were further informed that outcomes occurred independently of their actions and that the X indicated the absence of a reward rather than a penalty. Consequently, this phase involved reward and non-reward events rather than reward and loss, preventing the CS– from acquiring aversive properties.

On each trial, an empty slot-machine was displayed at the centre of the screen (ITI 500–1500 ms), followed by the presentation of one of the three CSs in the upper display for 2000ms. The corresponding outcome image then appeared in the lower display for 1000ms, while the CS remained visible in the upper window. No levers or point counter were presented during this phase, ensuring that participants learned the cue–outcome contingencies without performing instrumental responses.

A series of blocks, each composed of 45 intermixed trials (15 trials per CS), was repeated until the Pavlovian learning criterion was met. Cue presentation order was pseudo-randomized with the constraint that no CS could appear more than twice in succession. After each block, participants completed multiple-choice questions displayed on the screen to verify correct acquisition of the CS–outcome associations (e.g., “Which food did you earn with this stimulus?”, repeated for each CS). The learning criterion consisted of correctly identifying all CS–outcome associations after two consecutive blocks. If the criterion was reached, participants advanced to the next phase; otherwise, additional blocks were administered up to a maximum of eight. Thus, participants completed a minimum of two and a maximum of eight blocks. All participants met the learning criterion after the minimum of two blocks. After completing this phase, participants rated their subjective liking for each of the three CSs using a 9-point Likert scale (0–9) in response to the question, “How much do you like this stimulus?”. Compared with the baseline ratings, collected at the start of the task, this post-learning measure served to confirm the effectiveness of Pavlovian learning by assessing differential changes in stimulus valuation.

### Transfer phase

In the Transfer phase, participants performed again instrumental choices while Pavlovian cues were concurrently presented. This allowed us to assess the influence of previously acquired cue–outcome associations on instrumental responding, under varying volatility conditions. The structure and timing of the task mirrored the Instrumental learning phase, except for the addition of Pavlovian stimuli.

Each trial began with the presentation of an empty slot-machine (ITI 500–1500ms), followed by the appearance of the two levers on the left and right sides of the slot-machine and one Pavlovian cue (CS+₁, CS+₂, or CS–) displayed above it. The point counter was visible and updated trial by trial, as in the Instrumental phase.

Participants completed two blocks of 99 trials each, one for each volatility condition (low and high), resulting in a total of 198 trials. Within each block, each Pavlovian cue appeared on 33 trials, ensuring equal representation of CSs across the session. The reward contingencies followed the same 80/20 structure as in the Instrumental learning phase and each block contained the same number of reversals (one in the low- and five in the high-volatility context). Their exact trial positions, however, differed from the Instrumental phase, because reversals in the Transfer phase were additionally constrained so that the number of trials between consecutive reversals was a multiple of three, allowing each cue to appear equally often within each reversal period. Cue presentation order was pseudo-randomized under this constraint to avoid predictable sequences. The two blocks matched the volatility contexts experienced previously, presented in the same order (i.e., participants who began the Instrumental learning phase in the high volatility context also began the Transfer phase in that context). This was done to preserve the learned association between contextual cues (background colour) and volatility conditions, ensuring that the Transfer phase reinstated the same environmental context experienced during instrumental learning. Pavlovian cues were task-irrelevant; therefore, maximizing reward required responding solely based on instrumental contingencies, effectively ignoring the CSs.

#### Experimental Procedure

Participants were instructed to refrain from eating for at least 3 hours prior to the experiment to ensure a moderate state of hunger. Upon arrival at the laboratory, they provided written informed consent. The two rewarding outcomes were individually tailored for each participant by assessing subjective liking for a set of 10 food items (five savoury and five sweet). Foods were rated on a 5-point Likert scale ranging from 0 (*not at all*) to 5 (*very much*) and the two most highly and equally rated foods were selected as rewarding outcomes. Corresponding food images were then used as the reward stimuli throughout the task.

Before beginning the experimental session, participants completed an additional evaluation of the two selected foods using 9-point Likert scales (0–9) to assess both *general liking* (“How much do you usually enjoy eating it?”) and *current wanting* (“How much would you like to eat it now?”). Participants also rated their current hunger level on a 9-point scale to confirm comparable motivational states across individuals.

After completing these preliminary assessments, participants were seated comfortably in a quiet testing room, approximately 60 cm from a computer monitor and the two selected food rewards were placed in clear view on the table to maintain high motivational engagement throughout the task. Participants were informed that the total number of food pictures obtained during the task would determine the amount of food they would receive at the end of the experiment.

The task comprised three consecutive phases: an Instrumental learning phase, in which participants learned response–outcome associations under alternating low and high volatility contexts; a Pavlovian learning phase, in which participants learned cue–outcome associations in a neutral, non-volatile context; and a Transfer phase, in which Pavlovian influences on instrumental responding were tested under the same low and high volatility contexts. Participants were instructed to pay close attention to the on-screen instructions at the beginning of each phase. To ensure comprehension, a brief verbal summary of the task structure was provided by the experimenter before starting and a few example trials were administered for familiarization.

The entire experimental session lasted approximately 60 minutes. The experimenter remained in the room to monitor task performance and provide assistance if technical issues occurred. At the conclusion of the experiment, participants received at least one portion of each of the previously selected food rewards, with the total quantity proportional to their performance-based earnings.

#### Entropy-based metrics

To quantify trial-by-trial uncertainty in choice behaviour, we computed entropy-based metrics introduced by Trepka et al. (2021). These measures are grounded in Shannon’s information theory (Shannon, 1948), which formalizes uncertainty as the expected information content of a random variable. Entropy-based analyses have increasingly been used in reinforcement learning and decision-making research to capture variability or uncertainty in choice behaviour beyond mean performance levels (Trepka et al., 2021; Woo et al., 2023; Woo et al., 2025a).

In the present study, we focused on a metric that quantifies variability in the adopted choice strategy (stay vs. switch) conditional on the option chosen in the previous trial: the *Conditional Entropy of Option-Dependent Strategy* (EODS; Trepka et al., 2021). This measure can be decomposed into components reflecting strategic variability following a better or worse previous choice, respectively. Because our primary hypothesis concerns the disruption of optimal choice behaviour — that is, whether participants deviate from a stay-with-the-better-option strategy — we focused on EODS*B*, the conditional entropy of strategy given a previous better choice:

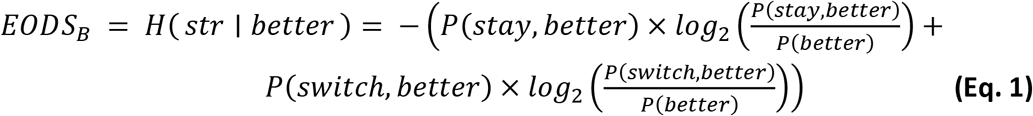

where *str* is strategy coded as stay (1) or switch (0). High EODS*B* values indicate greater unpredictability in responding after a correct choice. Because EODS*B* captures the consistency, but not the direction, of the choice strategy, we interpret it jointly with choice accuracy (P_better_): a concurrent rise in EODS*B* and fall in P_better_ indexes responding that is both less consistent and directed away from the better action: the pattern expected under Pavlovian interference.

#### Computational models

We fit a set of computational models to participants’ trial-by-trial choices during the Instrumental learning phase to test whether environmental volatility modulates learning dynamics.

##### i: Rescorla-Wagner with context-specific learning rates

The Rescorla-Wagner (RW) model (Sutton & Barto, 2018; Wagner & Rescorla, 1972) implements value learning via a simple delta rule. On each trial *t*, the agent first computes the reward prediction error (RPE):

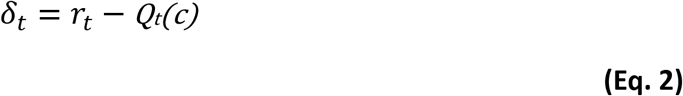

where *r_t_* ∈ {0,1} is the binary outcome and *Q_t_*(*c*) is the current expected value of the chosen option *c*. The expected value is then updated according to:

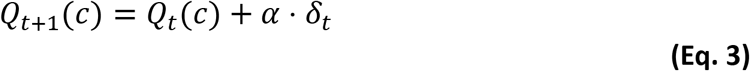

where *δ*_*t*_ represents the reward prediction error (RPE), and is calculated through the equation *r*_*t*_ − *Q*_*t*_(*c*), *r*_*t*_ ∈ {0,1} is the binary outcome and *α* ∈ [0,1] is the learning rate controlling the weight given to new information relative to prior beliefs (Sutton & Barto, 2018).

To test whether volatility modulates learning speed, we fit separate learning rates for low and high volatility blocks. The model thus had two learning rate parameters (α_Low_, α_High_) where blocks 1 and 3 correspond to one volatility condition and blocks 2 and 4 to the other (Behrens et al., 2007).

##### ii: Rescorla-Wagner with separate learning rates for positive and negative prediction errors

Extensive evidence suggests that learning from better-than-expected outcomes (positive RPEs) differs from learning from worse-than-expected outcomes (negative RPEs), with dissociable neural substrates in dopaminergic pathways (Daw et al., 2002; Frank et al., 2004). To capture this asymmetry, previous model extends the standard RW model by implementing separate learning rates for positive and negative prediction errors:

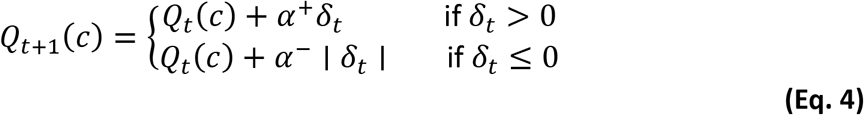

where *α*^+^ governs learning from rewards and *α*^−^ governs learning from omitted rewards. We fit separate *α*^+^and *α*^−^parameters for low and high volatility conditions (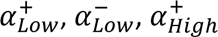, 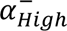), allowing us to test whether volatility differentially affects learning from positive versus negative outcomes.

##### iii: Pearce-Hall attention-weighted learning

The Pearce-Hall model (Pearce & Hall, 1980) proposes that learning rates are not fixed but dynamically adjust as a function of recent prediction error magnitude. This implements a form of attentional learning: agents allocate more attention—and thus learn faster—from stimuli that have recently generated large prediction errors. Following Roesch et al. (2012), we implemented the scaled version:

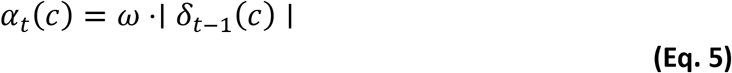

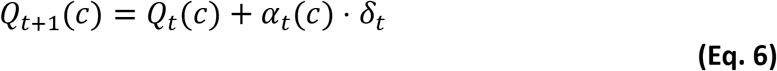

where *ω* is a scaling parameter controlling the sensitivity of learning rates to prediction errors. Crucially, the learning rate at trial t depends on the absolute prediction error from the previous trial on that option, implementing trial-by-trial adaptation. We fit separate ω parameters for low and high volatility blocks (*ω_Low_, ω_High_*).

This model bridges classical associative learning theory and modern reinforcement learning by providing a mechanistic account of how learning rates might adapt without requiring explicit representation of environmental statistics.

##### iv: Bayesian learning with discrete change-point detection

While preceding models adapt to volatility through fixed context-specific parameters, Bayesian models offer a complementary approach in which the rate of belief updating is derived endogenously from the observed sequence of outcome. By maintaining a full probability distribution over possible reward rates and updating this distribution via Bayes’ rule, these models provide a mechanistic account of how uncertainty is tracked and resolved trial by trial. (Behrens et al., 2007; Courville et al., 2006; Daw et al., 2005). We implemented a discretized Bayesian learner that represents uncertainty explicitly (Wilson & Niv, 2012). The model maintains a probability distribution *p*(*q*) over the reward rate *q* ∈ [0,1] for each option, discretized into 99 equally-spaced values. On each trial, the model updates beliefs via Bayes’ rule:

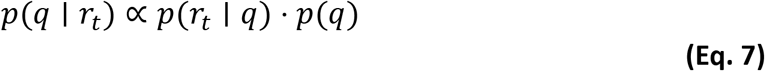

where the likelihood *p*(*r*_*t*_*| q)* is Bernoulli: *p*(*r*_*t*_=1 | *q*) = *q* and *p*(*r*_*t*_=0 | *q*) = *1−q*. The posterior distribution is computed for all 99 grid points and normalized. The expected value used for choice is the mean of the posterior distribution:

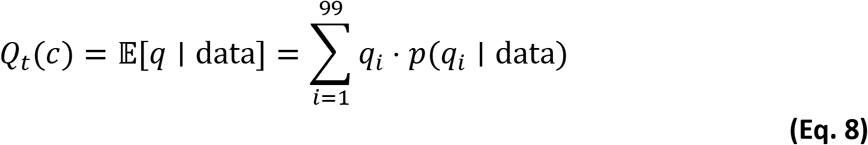

To model the possibility that reward contingencies can change (i.e., reversals), the posterior is mixed with a uniform distribution before the next trial:

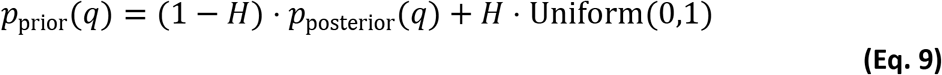

where *H* ∈ [0,1] is the hazard rate representing the agent’s prior belief about the probability of a change-point (reversal) occurring at any trial. High *H* values implement rapid “forgetting” of past data, equivalent to high learning rates in RW models, while low *H* values implement slow accumulation of evidence. Critically, we fit separate hazard rates for low and high volatility blocks (*H_Low_*, *H_High_*), testing whether participants adjust their change-point expectations to match environmental statistics.

This model represents the full Bayesian solution to learning in non-stationary environments but with a discrete rather than continuous representation (Wilson & Niv, 2012), making it computationally tractable while preserving key properties of optimal inference (Gershman, 2015).

##### v: Approximately Bayesian Delta-rule with Bernoulli likelihood

While previous model implements change-point detection via fixed hazard rates, this (Nassar et al., 2010, 2012) computes the probability of a change-point dynamically on each trial by comparing the likelihood of observed outcomes under two hypotheses: (1) reward contingencies have remained stable versus (2) a change-point has occurred. This trial-by-trial inference allows for more flexible adaptation.

The model maintains a point estimate of the reward rate *r̂_t_*(*c*) and uncertainty 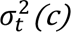 for each option. Upon observing outcome *y*_*t*_, the model computes the change-point probability Ω_*t*_ using Bayes’ rule:

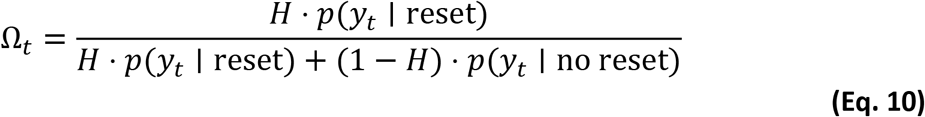

where *H* is the prior probability of a change-point (hazard rate), *p*(**y**_*t*_ | reset) assumes the reward rate has reset to a uniform prior (*r̂* = 0.5) and *p*(**y*_*t*_* | no reset) uses the current estimate. For binary outcomes, we use Bernoulli likelihoods:

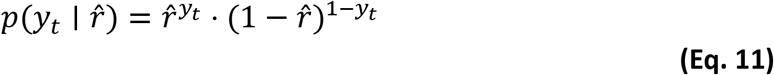

The learning rate is then computed as a mixture of complete reset (Ω_*t*_ → 1) and incremental Kalman-like update (Mathys et al., 2014; Piray & Daw, 2020, 2021):

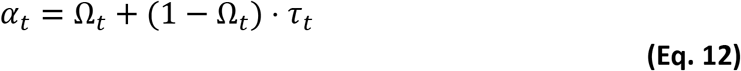

where 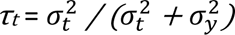 is the Kalman gain, with 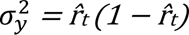 representing the variance of a Bernoulli distribution and *σ*^2^ is a noise scaling parameter. The reward rate estimate is updated as:

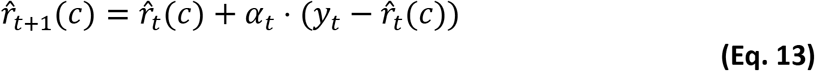

Uncertainty is also updated, increasing after putative change-points and decreasing as evidence accumulates:

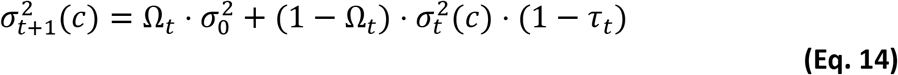

where 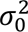 = 0.25 is the prior uncertainty after a reset, corresponding to the maximum variance of a Bernoulli distribution (i.e., Var = p(1−p) at p = 0.5), which encodes maximal uncertainty about the reward rate following a putative change-point. We fit separate hazard rates (*H_Low_*, *H_High_*) and noise parameters (**σ*_Low_*, **σ*_High_*) for each volatility condition. This model predicts that in high volatility, Ω_*t*_will be elevated on average, producing higher effective learning rates.

This model has been widely applied to characterize human learning in volatile and non-stationary environments, as the trial-by-trial change-point inference provides a flexible mechanism for adapting to abrupt shifts in reward contingencies (McGuire et al., 2014; Nassar et al., 2010, 2012). For this model, the value entering the choice rule corresponds to the estimated reward rate of each option: that is, *Q_t_*(*c*) = *r̂_t_*(*c*).

All models used a softmax decision rule to map values to choice probabilities:

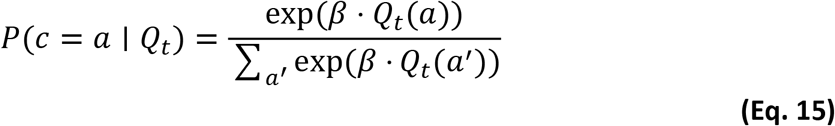

where *β* is the inverse temperature parameter controlling the stochasticity of choices. Higher *β* values produce more deterministic choices (exploitation), while lower values produce more random exploration. We fit separate inverse temperature parameters for low and high volatility blocks (*β_Low_*, *β_High_*) to allow for the possibility that choice stochasticity varies with volatility.

For each model, free parameters were estimated using a maximum a posteriori estimation (Gershman, 2016) using the mfit toolbox (https://github.com/sjgershm/mfit), which performs the optimization via *fmincon* (MATLAB Optimization Toolbox) to find the parameter values that maximized the posterior probability of the observed choices. To prevent local minima, we initialized each optimization with 25 random starting points sampled from uniform distributions over plausible parameter ranges (inverse temperatures: [0, 50]; learning rates and hazard rates: [0, 1]; noise parameters: [0, 1]). For the very first block of the session only, value estimates for the two options (Q) were initialized at 0.5. For the blocks that followed switches, initial values were set equal to Q values from the last trial of the previous block, given by its best-fit parameters. Because all participants met the learning criterion after the minimum of four blocks (see Instrumental learning phase), models were fitted to 200 trials per volatility condition (two 100-trial blocks per condition) for every participant. Models were fitted separately to low- and high-volatility blocks to test the hypothesis that high volatility increases learning rates.

Goodness-of-fit was assessed using the Bayesian Information Criterion (BIC), to identify the model that best captured the behaviour. Additionally, we performed a parameter recovery analysis to verify that the fitting procedure can reliably identify the parameters that generated the data. For each participant, we simulated a synthetic dataset using the individually fitted parameters of the winning model and re-fitted the model to the simulated data (Palminteri et al., 2017; Wilson & Collins, 2019). The correspondence between generating and recovered parameters was quantified by Pearson correlations across participants, with 95% confidence intervals estimated via non-parametric bootstrap (10,000 resamples).

Then, to validate the winning model’s ability to capture the main aspects of the data, we compared the observed instrumental response rates with those generated from 100 simulated datasets. These simulations were conducted using the participants’ fitted parameters and matched the sample size of the empirical data.

Finally, we used the estimated model parameters from the best-fitting model to investigate differences in the learning dynamics across volatility contexts.

#### Statistical analysis

All data were processed offline using custom-made MATLAB scripts (The MathWorks, Inc., Natick, MA, USA). Statistical analyses were carried out using JASP 0.19 (Love et al., 2019). As preregistered, we adopted a dual inferential approach combining null hypothesis significance testing (NHST) and Bayesian inference. Normality and sphericity assumptions were visually inspected through Q–Q plots and by verifying that skewness and kurtosis values were < |2| for all variables (Hopkins & Weeks, 1990). When assumptions for parametric tests were not met, Bayesian non-parametric tests were adopted (e.g., Bayesian Mann–Whitney), with Kendall’s W reported as an effect size.

For the Instrumental learning phase, behaviour was quantified as the percentage of choices of the currently better option (P_better_), computed separately for the Low- and High-volatility contexts and for each session, providing context-specific estimates of performance. In addition to P_better_, we ran the same context- and session-wise analyses using EODS*_B_* (the conditional entropy of choices given that the previous trial featured the better action). EODS*_B_* was chosen for congruence with the main analyses (i.e., reported relative to the better action) and provides a complementary measure of choice uncertainty. In the Transfer phase, performance was quantified not only as P_better_ and EODS*_B_* computed separately for each volatility context, but also as a function of the Pavlovian cues presented on each trial, allowing us to assess how previously acquired Pavlovian associations influenced instrumental responding. To characterize transfer effects independently of the reversal schedule, Pavlovian cues were additionally recoded according to their relation to the current instrumental contingency: CS_b_, predicting the outcome associated with the currently better action (i.e., the *better* CS); CS_w_, predicting the outcome associated with the currently worse action (i.e., the *worse* CS); and CS–, predicting the non-reward outcome. For the Transfer phase, EODS*_B_* was computed separately for each Pavlovian cue as recoded according to the current instrumental contingency (CS_b_, CS_w_, CS–), by selecting trials in which the immediately preceding choice had been the better action. This allowed us to index cue-specific choice entropy conditional on a prior better response.

NHST was performed using a significance threshold of *p* < .05. Effect sizes are reported as partial η² for ANOVAs (Lakens, 2013) and Cohen’s *d* for paired-samples t-tests. Bayesian inference was used to provide graded evidence for or against effects of interest. For all ANOVAs and t-tests, we report the Bayes Factor (BF₁₀), expressing the likelihood of the data under the alternative hypothesis relative to the null hypothesis (Kruschke, 2021). All Bayesian tests used JASP’s default prior (Cauchy distribution with scale r = 0.707). Interpretation of Bayes Factors follows established guidelines (Andraszewicz et al., 2015; Lee & Wagenmakers, 2014): BF₁₀ < 1/3 was taken as moderate evidence for H₀, BF₁₀ > 3 as moderate evidence for H₁ and values between 1/3 and 3 as inconclusive. Stronger thresholds follow Lee and Wagenmakers (2014), with BF₁₀ > 100 indicating extreme evidence for H₁ and BF₁₀ < 1/100 extreme evidence for H₀.

In addition, Transfer phase differences between each Pavlovian cue condition (CS_b_, CS_w_) and the neutral control (CS–) were quantified via estimation statistics (Cumming, 2014; Ho et al., 2019). For each participant, paired differences (CS_b_ − CS–; CS_w_ − CS–) were computed separately for each volatility context and, in Experiment 2, for each feedback condition (Probe vs. Normal). Estimation plots show individual paired differences together with the group mean difference (Δ_mean_) and its 95% bias-corrected accelerated (BCa) bootstrap confidence interval (CI), based on 5000 bootstrap resamples (Degni et al., 2024). Inference was based on the magnitude of Δ_mean_ and the precision of its estimate (i.e., CI width): CIs including zero were interpreted as indicating no reliable evidence of a cue-driven effect, whereas CIs excluding zero were interpreted as indicating reliable evidence, with the strength of the effect further qualified by its magnitude and the width of the CI (Calin-Jageman & Cumming, 2019).

## Results

### Outcomes liking and wanting

Participants showed comparable liking (t_49_ = – 0.562; p = 0.577; d = – 0.079; BF₁₀ = 0.179) and wanting ratings (t_49_ = 0.586; p = 0.561; d = 0.083 BF₁₀ = 0.181) for the two rewarding outcomes at the start of the task, confirming the effectiveness of the reward-selection procedure.

### Acquisition of Pavlovian learning

All participants (N = 50; 100%) reached the learning criterion, correctly answering the CS–outcome association question after the minimum two blocks required. Subsequently, we analysed CS-liking ratings using a 2 × 3 repeated-measures ANOVA with Time (pre vs. post) and CS (CS+_1_, CS+_2_, CS–) as within-subject factors. The analysis revealed a significant CS × Time interaction (F_2,98_ = 29.695; p < 0.001; η_p_² = 0.377; BF₁₀ = 9.710×10^8^), as well as significant main effects of CS (F_2,98_ = 4.831; p = 0.010; η_p_² = 0.090; BF₁₀ = 3.997) and Time (F_1,49_ = 15.594; p < 0.001; η_p_² = 0.241; BF₁₀ = 21.325) (**Fig. 2a**). Inspection of the means and 95% CI indicated that, prior to Pavlovian learning, the three CSs were rated comparably. Following learning, both CS+_1_ and CS+_2_ were liked more than CS– and liking increased from pre- to post-learning for the CS+ stimuli, whereas it decreased for the CS– stimulus. This pattern reflects successful acquisition of the Pavlovian cue–outcome associations.

**Fig. 1.**
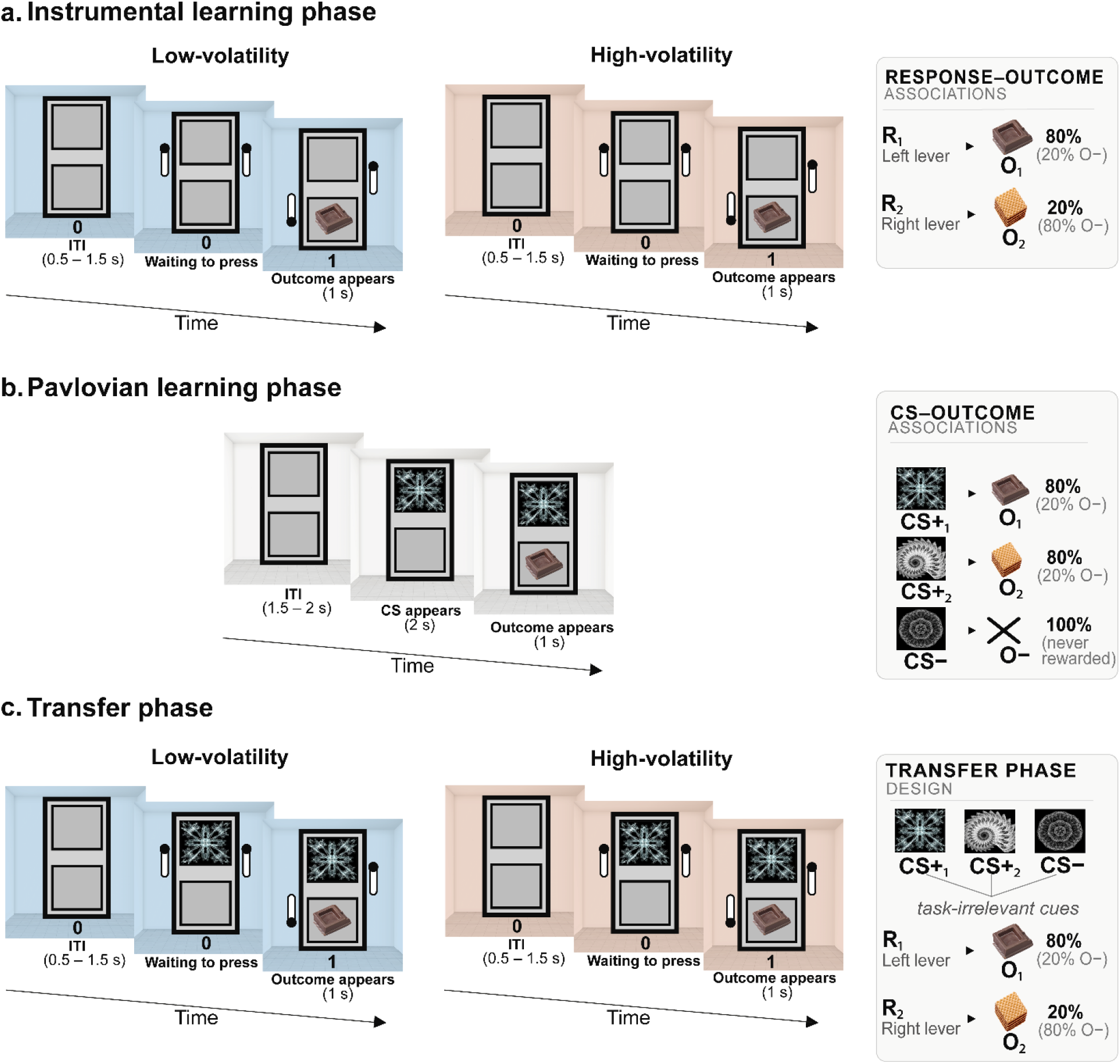
Illustration of the Pavlovian-to-Instrumental Transfer task with volatility manipulation. **(a)** Instrumental learning phase. On each trial, an empty slot-machine appeared (ITI 500–1500 ms), followed by two levers (R_1_, R_2_) presented on either side; responses were followed by one of two food outcomes (O_1_, O_2_) delivered according to an 80/20 probabilistic schedule. A coloured background room (light-blue or orange) surrounding the slot-machine signalled the low- or high-volatility context throughout the block. **(b)** Pavlovian learning phase. Participants passively observed the association between three conditioned stimuli (CS+_1_, CS+_2_, CS−), shown above the slot-machine and their corresponding outcomes (O_1_, O_2_, or no reward), in the absence of levers or instrumental responding. **(c)** Transfer phase. The instrumental display was reinstated (levers and point counter visible) together with one of the three Pavlovian cues, presented above the slot-machine, allowing cue-driven and instrumentally optimal responses to be measured concurrently under the low- and high-volatility contexts established during instrumental learning.

**Fig. 2.**
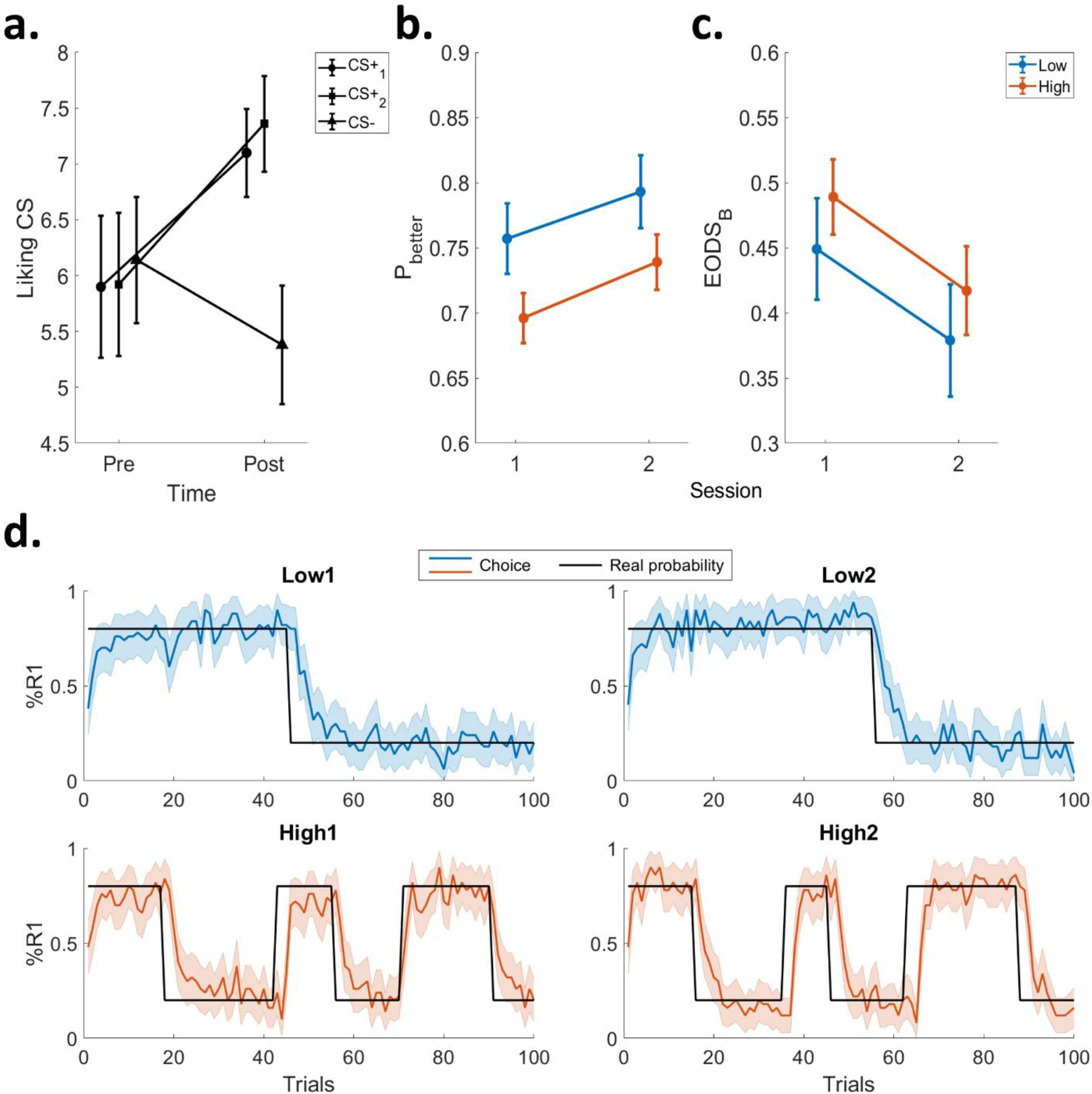
Pavlovian learning and instrumental performance across volatility contexts. **(a)** Cue–outcome learning during Pavlovian conditioning. CS-liking ratings are shown before and after conditioning for the two reward-predictive cues (CS+_1_, CS+_2_) and the non-reward cue (CS–). Bars represent group means with 95% confidence intervals. Before learning, cues were rated similarly; after learning, liking increased for the reward-predictive cues and decreased for CS–, consistent with successful acquisition of Pavlovian associations. **(b)** Choice accuracy (*P*_better_) during instrumental learning as a function of Session (1 vs. 2) and Volatility (Low vs. High). Bars represent group means with 95% confidence intervals. Participants improved from Session 1 to Session 2 and performance was higher under Low volatility, indicating more robust exploitation of the optimal action when environmental statistics were stable. **(c)** Choice uncertainty (EODS*_B_*): conditional entropy of the current choice given that the preceding trial featured better action. Lower entropy in Low volatility reflects more consistent engagement with the better-choice strategy relative to High volatility. Bars represent group means with 95% confidence intervals. **(d)** Trial-by-trial choice behaviour in the instrumental phase. The y-axis shows %R1 (percentage of selections of the initially better action), while the x-axis represents trials number. Blue lines display low-volatility performance (top panels), orange lines high-volatility performance (bottom panels). Shaded bands indicate 95% CI across participants and the black line shows the true reward probability of R1.

### Effects of environmental volatility on instrumental choice behaviour

In the instrumental learning phase, all participants met the learning criterion after the minimum two sessions, as evidenced by correct responses to the action–outcome association question. To assess whether environmental volatility modulated instrumental performance (indexed by *P*better i.e., the probability of selecting the currently optimal action), we conducted a 2 × 2 repeated-measures ANOVA with Session (1 vs. 2) and Volatility (Low vs. High) as within-subject factors. The analysis revealed a significant main effect of Session (F_1,49_ = 23.565; p < 0.001; η_p_² = 0.325; BF_10_ = 168.742), with higher *P*better in Session 2 than Session 1, indicating overall improvement in instrumental learning with training. A significant main effect of Volatility also emerged (F_1,49_ = 33.346; p < 0.001; η_p_² = 0.405; BF_10_ = 1.451×10^4^), showing better performance in the Low-volatility context compared to the High-volatility context (**Fig. 2b**). The interaction between the two factors was not significant (F_1,49_ = 0.151; p = 0.699; η_p_² = 0.003; BF_10_ = 0.216). These results confirms that instrumental accuracy differed across volatility levels, with reduced performance under high volatility, consistent with previous reports of diminished choice precision in unstable environments (Farashahi et al., 2019; Soltani et al., 2021).

We ran the same 2 × 2 repeated-measures ANOVA on EODS*_B_* (the conditional entropy of choices when the previous trial featured the better action) to test whether volatility also modulated choice uncertainty. This ANOVA produced analogous effects to those observed for *P*better: a main effect of Session (F_1,49_ = 33.725; p <0.001; η_p_² = 0.408; BF₁₀ = 11.196×10^3^), a main effect of Volatility (F_1,49_ = 6.114; p = 0.017; η_p_² = 0.111; BF₁₀ = 3.104) and a non-significant Session × Volatility interaction (F_1,49_ = 0.010; p = 0.922; η_p_² = 1.958×10^-4^; BF₁₀ = 0.201) (**Fig. 2c**). In line with the behavioural accuracy results, EODS*_B_* decreased from Session 1 to Session 2, reflecting more consistent better-choice behaviour as learning progressed and was lower in the Low- than in the High-volatility (indicating increased choice uncertainty).

A complementary inspection of trial-by-trial choice behaviour (**Fig. 2d**) further illustrates how participants performed across sessions and volatility contexts. Performance remained consistently above chance in both conditions (M*Low*=0.775; SD*Low* = 0.080; M*High* = 0.718; SD*High* = 0.059) and accuracy increased from the first to the second session (M*Low1* = 0.757, SD*Low1* = 0.092; M*Low2* = 0.793, SD*Low2* = 0.096; M*High1* = 0.696, SD*High1* = 0.067; M*High2* = 0.739, SD*High2* = 0.072). These patterns confirm that participants were not responding randomly but successfully adapted their choices to the structure of the task.

### Computational modelling reveals adaptive learning rate modulation in volatile environments

To confirm that participants differentiated between volatility contexts at the computational level, as predicted by theoretical accounts of learning rate adaptation (Behrens et al., 2007; Massi et al., 2018), we fitted trial-by-trial choice data using five computational models grounded in different theoretical frameworks (see Methods for details). The models included: (1) a Rescorla-Wagner (RW) model with separate learning rates and inverse temperatures for low and high volatility contexts; (2) a RW model with separate learning rates for positive and negative prediction errors (RW +/-) for both low and high volatility contexts; (3) a Pearce-Hall model with attention-weighted learning; (4) a Bayesian learning model with explicit uncertainty estimation; and (5) an approximate Bayesian-Delta rule with change-point detection model. To determine the best-fitting model, we used the BIC as measure of goodness-of-fit (Wilson & Collins, 2019). We found that the best model was the Rescorla-Wagner, with separate learning rates (α) and inverse temperatures (β) for low and high volatility blocks (mean BIC = 351.42; see Supplementary Table S1 for BIC values and estimated parameter distributions of all models) (**Fig. 3a**). This model outperformed all alternative models, indicating that participants’ behaviour was best characterized by context-dependent modulation of both learning rates and decision noise. Crucially, we performed model validation to ensure that the best-fitting model captures the main aspects of the behavioural data. For model validation, we simulated 100 datasets using the fitted individual parameters and compared key behavioural signatures between real and simulated datasets. First, we examined overall choice accuracy (*P*better) across both Session (1 and 2) and Volatility (Low and High) as in **Fig.2b**. The model successfully reproduced the observed pattern of performance, capturing both the main effect of volatility and session effects (**Fig. 3b**). Additionally, we examined participants’ ability to rapidly adjust their choices following reward contingency reversals, as this behavioural signature is particularly diagnostic of learning mechanisms. We compared the time course of choice adjustment in a ±10 trial window centred on reversal points. The model accurately reproduced the characteristic pattern of adjustment around reversals (**Fig. 3c**), including: (i) the initial drop in performance immediately following a reversal; (ii) the subsequent recovery trajectory over the following 10 trials; and (iii) the differential adjustment dynamics between volatility conditions, with faster recovery in high compared to low volatility contexts. Overall, these validation analyses demonstrate that the best-fitting model captures both overall choice patterns and fine-grained trial-by-trial dynamics of adaptive learning behaviour.

**Fig. 3.**
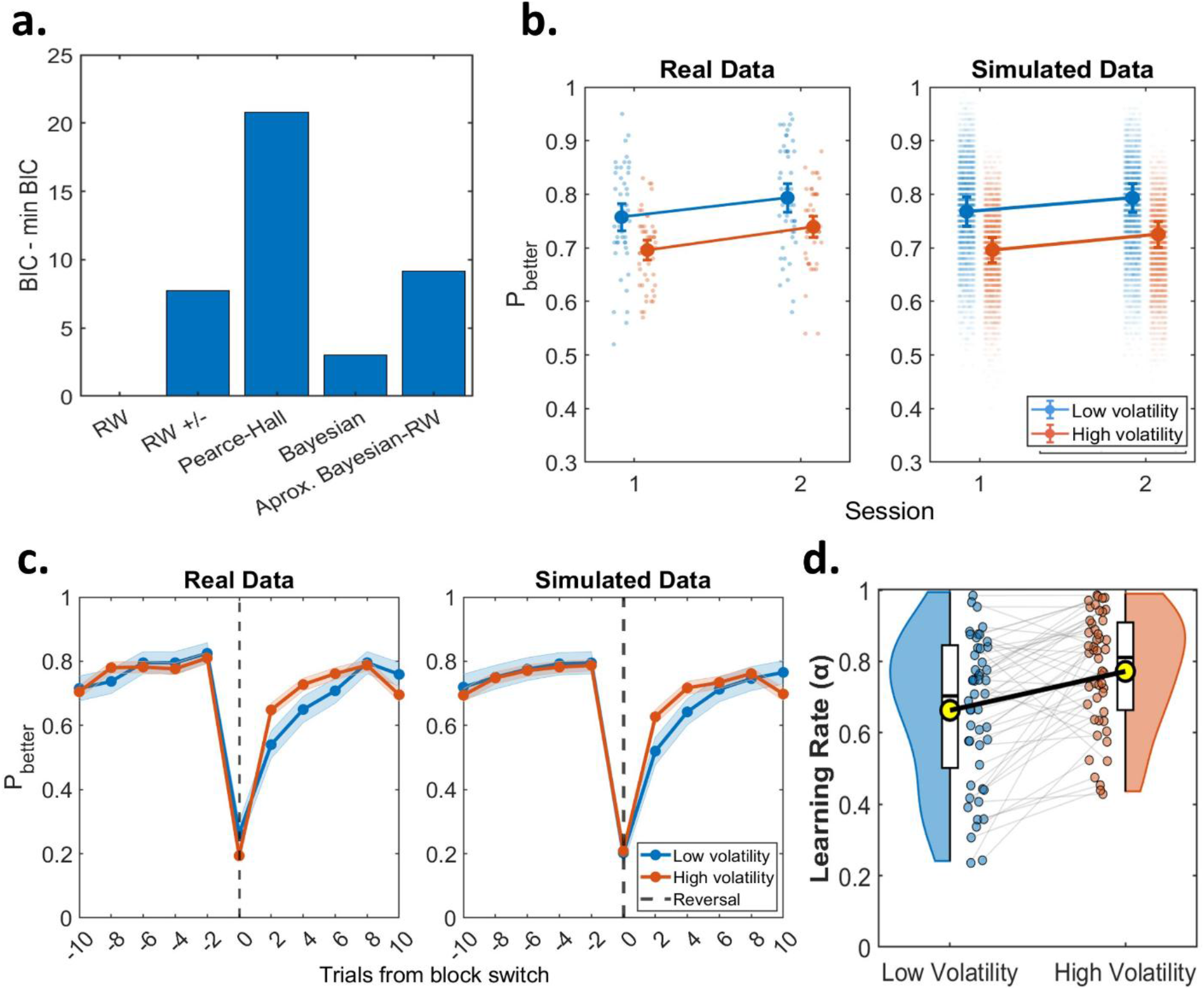
Goodness-of-fit for various computational models, model validation and model estimated parameters. **(a)** Bayesian information criteria for different reinforcement learning models. **(b)** Model validation: choice accuracy (*P*_better)_. Comparison between real (left) and simulated (right) data across the 2×2 experimental design (Session × Volatility). Individual data points represent single subjects’ mean performance in each condition; Larger dots with error bars represent group means ± 95% CI. **(c)** Model validation: adjustment to reward contingency reversals. Time course of choice accuracy (*P*_better_) in a ±10 trial window centred on reversal points. The model accurately captures the characteristic pattern of behavioural adjustment. Critically, the model reproduces the differential adjustment dynamics between volatility conditions, with faster recovery in high (orange) compared to low (blue) volatility contexts. Left panel: real data; right panel: simulated data. Coloured dots with shaded regions represent time point means ± 95% CI. Vertical dotted line represents contingencies reversal. **(d)** Estimated learning rate parameters (α) from the best-fitting Rescorla–Wagner model for Low and High volatility contexts. Half-violin and box plots show the distribution of individual parameter estimates; boxes indicate the interquartile range with the median; grey lines connect the same participant across conditions; yellow dots connected by a black line indicate group means.

To test our hypothesis regarding the adaptive modulation of learning rates as a function of environmental volatility, we performed a paired-samples *t*-test across participants to compare the estimated learning rate parameters between low and high volatility conditions (α_Low_, α_High_). The results shown a strongly significant difference in the learning rates across volatility contexts (t_49_ = – 3.109; p = 0.003; d_Cohen_ = – 0.440; BF_10_ = 10.361). Visual inspection of the results (**Fig. 3d**) shows that learning rates were higher in high volatility contexts compared to low volatility contexts, reflecting that participants dynamically adjusted the rate at which they weighted new information in response to environmental instability, a pattern that aligns with established findings in both human (Behrens et al., 2007) and non-human primate (Massi et al., 2018) studies. To verify that this adaptation was specific to the learning rate rather than reflecting a broader change in choice stochasticity, we performed an analogous paired-samples *t*-test on the estimated inverse temperature parameters (β_Low_, β_High_). In contrast to the learning rate, the inverse temperature did not differ between volatility contexts (*t*₄₉ = – 0.737; *p* = 0.447; *d* = – 0.109; BF₁₀ = 0.203, moderate evidence for H₀), indicating that volatility selectively modulated the speed of value updating while leaving choice consistency unchanged (Supplementary Fig. S2a). Finally, a parameter recovery analysis confirmed that the fitting procedure reliably identified the generative parameters of the winning model: correlations between simulated and recovered values were strong for all free parameters (all *r* > 0.79; 95% bootstrap CIs in Supplementary Fig. S3a), supporting the interpretability of the individual estimates.

To assess whether the volatility-dependent modulation of learning parameters observed for the winning model generalized across alternative decision architectures, we additionally compared, for all five candidate models, the free parameter(s) governing context-specific value updating and the inverse-temperature parameter between Low and High volatility (Supplementary Tables S3, S4). With the exception of the approximate Bayesian delta-rule model (Nassar et al., 2010, 2012), all models showed a reliable difference in their update-rule parameter(s) between volatility contexts, consistent with the pattern observed for the winning Rescorla–Wagner model; within the model with asymmetric learning rates, this reliable difference was specific to the negative-outcome learning rate (α⁻), whereas the positive-outcome learning rate (α⁺) did not differ between contexts. Inverse-temperature parameters did not differ between contexts in any model, mirroring the results reported above for the winning model (Supplementary Fig. S2a).

### Volatility shapes the expression of Pavlovian bias in the Transfer phase

To test our preregistered hypothesis that Pavlovian bias in the transfer phase would be modulated by the volatility learned during the instrumental learning, we analyzed transfer performance both in terms of choice accuracy (*P*better) and choice uncertainty (EODS*_B_*). Pavlovian stimuli were recoded relative to the current instrumental contingency as CS_b_ (cue congruent with the currently better action), CS_w_ (cue congruent with the currently worse action, i.e., conflict trials) and CS– (neutral control cue).

A 3 (CS: CS_b_, CS_w_, CS–) × 2 (Volatility: Low, High) repeated-measures ANOVA on *P*better (**Fig. 4a**) revealed a main effect of CS (F_2,98_ = 24.357; p <0.001; η_p_² = 0.332; BF₁₀ = 5.052×10^6^), a main effect of Volatility (F_1,49_ = 6.114; p = 0.017; η_p_² = 0.595; BF₁₀ = 8.741×10^6^) and a significant CS × Volatility interaction (F_2,98_ = 71.923; p <0.001; η_p_² =0.209; BF₁₀ = 1.061 ×10^4^). **Fig. 4a** (*P*better with 95% confidence intervals) shows a summary of such results. First, performance in the Low-volatility condition is consistently higher than in the High-volatility condition across cue types, mirroring the pattern observed during the instrumental phase. Second, CS_b_ produces a robust facilitation relative to CS−: estimation statistics (Supplementary Fig. S4) confirmed a reliably positive paired mean difference in both volatility contexts (Low: Δ_mean_ = 0.046, 95% CI [0.014, 0.079]; High: Δ_mean_ = 0.044, 95% CI [0.012, 0.076]), indicating that cues congruent with the optimal action supported the optimal strategy irrespective of volatility. Crucially, only the conflict cue (CS_w_) behaved differently depending on prior volatility: the CS_w_–CS− difference was centred near zero in the Low-volatility context (Δ_mean_ = −0.023, 95% CI [−0.051, 0.007]), indicating no reliable cue-driven disruption, whereas in the High-volatility context it was reliably negative (Δ_mean_ = −0.134, 95% CI [−0.183, −0.091]), consistent with the marked drop in *P*_better_ relative to CS− that drove the observed interaction. These findings indicate stronger Pavlovian bias (i.e., cue-driven deviation from the better action) during high-volatility context, consistent with our preregistered expectation.

**Fig. 4.**
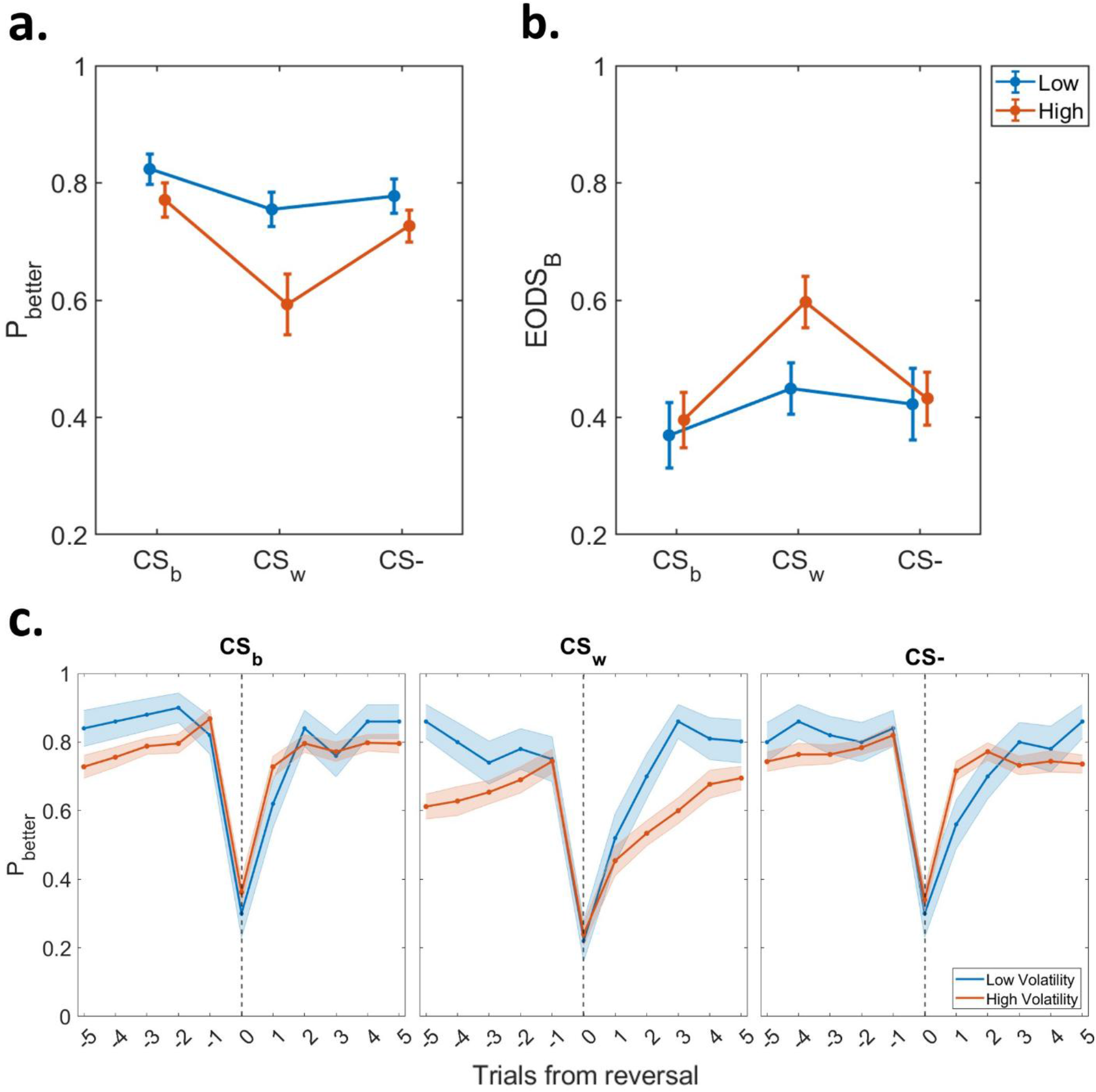
Volatility-dependent Pavlovian modulation of instrumental performance in the Transfer phase. **(a)** Choice accuracy (*P*_better_) as a function of Pavlovian cue type and volatility context (Low vs. High). Bars show group means and 95% confidence intervals. A main effect of Volatility indicates overall reduced accuracy following high-volatility learning. CS_b_ (cue congruent with the currently optimal action) facilitates optimal responding across contexts, yielding the highest *P*_better_. Critically, CS_w_ (cue congruent with the currently worse action) selectively disrupts performance in the High-volatility condition, producing the observed CS × Volatility interaction. **(b)** Choice uncertainty (EODS*_B_*: conditional entropy given a prior better response) for the same conditions. Lower entropy indicates more consistent better-choice behaviour. EODS*_B_* is generally reduced in Low volatility relative to High volatility and is minimized for CS_b_ across contexts. Crucially, CS_w_ markedly increases choice uncertainty only after high-volatility learning, mirroring the results in panel (a). Error bars represent 95% confidence intervals. **(c)**Reversal-locked *P*_better_ across a ±5-trial window centred on each contingency reversal, by CS condition and volatility context. Shaded regions represent 95% confidence intervals. In the CS_b_ and CS− conditions, post-reversal recovery is faster under high volatility; in the CS_w_ condition, the pattern reverses, indicating that conflicting Pavlovian cues anchor responding to the no-longer-optimal action and counteract the otherwise faster instrumental updating.

We complemented the accuracy analysis with a 3 × 2 repeated-measures ANOVA on EODS*_B_* (choice entropy conditional on a prior better response). The EODS*_B_* analysis paralleled the *P*better results, showing a main effect of CS (F_2,98_ = 15.856; p <0.001; η_p_² = 0.241; BF₁₀ = 10.393×10^3^), a main effect of Volatility (F_1,49_ = 14.198; p <0.001; η_p_² = 0.221; BF₁₀ = 11.302) and a significant CS × Volatility interaction (F_2,98_ = 6.080; p = 0.003; η_p_² = 0.108; BF₁₀ = 55.482).

The pattern in **Fig. 4b** closely mirrors the previous results: EODS*_B_* is reduced under low relative to high volatility, indicating more consistent better-choice behaviour in the stable context; CS_b_ yields the lowest entropy values across contexts and CS_w_ selectively increases entropy under high volatility relative to CS–, while overlapping with CS– in low, this indicates that in a volatile environment, conflicting cues induce greater uncertainty regarding the better choice.

To examine the temporal dynamics of this effect, we computed reversal-locked *P*better trajectories across a window of five trials before and after each contingency reversal, separately for each CS condition and volatility context (**Fig. 4c**). In the CS_b_ and CS− conditions, high-volatility participants showed faster post-reversal recovery, consistent with the elevated learning rates documented above. Critically, this pattern was reversed in the CS_w_ condition: post-reversal adjustment was slower under high volatility, with accuracy remaining below the low-volatility trajectory for several trials, indicating that the conflicting Pavlovian cue anchored responding to the no-longer-optimal action, counteracting the faster instrumental updating otherwise expected under high volatility. This dissociation rules out a general performance deficit and provides trial-by-trial evidence that volatility selectively amplifies Pavlovian interference when cue-driven and goal-directed signals conflict.

Taken together, the transfer results confirm our hypothesis: learning under high volatility increases the influence of Pavlovian cues on instrumental choice, producing both a reduction in optimal responding and an increase in cue-specific better-choice uncertainty (EODS*_B_*) when Pavlovian cues predict the currently worse action (CS_w_ trials).

## Experiment 2

Experiment 1 showed that high volatility shifts the balance of control toward the Pavlovian system when outcome feedback is available. Yet this shift might have been sustained by the feedback itself, rather than by the prior experience of volatility. Experiment 2 was designed to disentangle these two possibilities by measuring this shift without outcome feedback. Removing feedback entirely, however, would also prevent participants from tracking the current volatility context, the very factor whose lasting influence we aimed to isolate. We therefore restructured the Transfer phase to alternate brief Probe segments, run without outcome feedback, with Normal segments, run with outcome feedback, which sustained participants’ sense of the prevailing volatility. We first describe this design, then ask whether the volatility-dependent shift observed in Experiment 1 persists without outcome feedback.

### Methods

#### Participants

A different independent sample of twenty-eight healthy adult volunteers (age ≥ 18 years) (16 females, mean age = 23.68; sd = 3.41 years; mean education = 16.25; sd = 1.77 years) was recruited for the experiment. Participants were naïve to the purposes of the experiment and provided written informed consent prior to participation. The study was conducted in accordance with institutional guidelines and the 1964 Declaration of Helsinki and was approved by the Bioethics Committee of the University of Bologna.

The target sample size was determined a priori based on a preregistered power analysis (<u>10.17605/OSF.IO/Q2CMN</u>) conducted using MorePower 6.0 (Campbell & Thompson, 2012) for the planned 3 (CS: CS+1, CS+2, CS–) × 2 (Volatility: Low, High) repeated-measures ANOVA on transfer-phase data. The analysis assumed α = 0.05 (two-tailed), an effect size of η_p_² = 0.209 for the CS × Volatility interaction (estimated from the previously study using the same experimental design) and a desired power of 0.90. This analysis indicated that a sample size of 28 participants would be sufficient to detect the effect of interest and was expected to yield strong Bayesian evidence in favour of the alternative hypothesis (expected BF₁₀ = 12.67; (M. D. Lee & Wagenmakers, 2014). All participants were included in the analysis.

#### Experimental Paradigm

The task was identical to Experiment 1 except for the Transfer phase. The Transfer phase was redesigned in Experiment 2 to isolate Pavlovian influences on instrumental choice preventing concurrent reinforcement-driven updating of action or cue values. Specifically, we implemented this phase under nominal extinction (Badioli et al., 2024; Cartoni et al., 2015; Degni et al., 2022), i.e. participants were instructed that they were still earning food but, since the lower display of the slot machine was malfunctioning, they would not be able to see the outcomes and the total number of points earned. By removing immediate feedback while keeping the contextual volatility signals visible, this design tests whether prior exposure to high versus low volatility alone (rather than ongoing reinforcement) increases the expression of Pavlovian bias on choice, in line with our preregistered hypotheses.

Each Transfer block corresponded to one volatility context and comprised 102 trials. Blocks began with a 12-trial Anchor segment (no cues; feedback and point counter visible; identical to the Instrumental learning phase) to reinforce the current volatility context. The Anchor was followed by three Probe–Normal cycles; each cycle included a 12-trial (4 trials for each CSs) Probe segment (one Pavlovian cue presented on each trial alongside the two action options; feedback and the point counter withheld) immediately followed by an 18-trial (6 trials for each CSs) Normal segment (one Pavlovian cue presented on each trial alongside the two action options; feedback and point counter reinstated). Thus, each 102-trial block comprised 12 Anchor trials, 54 Normal trials (18 per CSs) and 36 Probe trials (12 per CSs).

Reversals were restricted to Normal segments: under low volatility a single reversal occurred approximately midway through the block (specifically in the second Normal segment), whereas under high volatility one reversal occurred late in the Anchor (≈trial 8) and four additional reversals were distributed approximately evenly across the Normal segments (≈ every 9–18 trials), yielding five reversals per high volatility block (mirroring the number of reversals during the Transfer phase in Experiment 1). To ensure balanced cue exposure across reversal periods, mini-blocks between consecutive reversals comprised a number of trials that was a multiple of three and cue order was pseudo-randomized under this constraint (as in Experiment 1).

This procedure provides repeated, well-controlled Probe segments in which to measure Pavlovian modulation of instrumental choice while preserving participants’ ability to track contextual volatility.

Procedure and statistical analysis approaches were otherwise identical to those used in Experiment 1.

### Results

#### Outcomes liking and wanting

Participants exhibited comparable liking (t_49_ = - 0.398; p = 0.693; d = - 0.075; BF₁₀ = 0.216) and wanting ratings (t_27_ = - 0.386; p = 0.702; d = - 0.072; BF₁₀ = 0.215) for the two rewarding outcomes at task onset, indicating that the reward-selection procedure successfully equated their subjective value.

#### Acquisition of Pavlovian learning

All participants (N = 28; 100%) met the learning criterion, correctly identifying the CS–outcome associations after the minimum two blocks required. CS-liking ratings were then analyzed with a 2 × 3 repeated-measures ANOVA with Time (pre vs. post) and CS (CS+_1_, CS+_2_, CS–) as within-subject factors. The analysis revealed a significant CS × Time interaction (F_2,54_ = 9.224; p < 0.001; η_p_² = 0.255; BF₁₀ = 4.098×10^2^) and a main effect of Time (F_1,27_ = 7.722; p = 0.010; η_p_² = 0.222; BF₁₀ = 2.312), whereas the main effect of CS was not significant (F_2,54_ = 2.060; p = 0.137; η_p_² = 0.071; BF₁₀ = 0.569) (**Fig. 5a**). Inspection of the means showed that the three CSs were rated similarly prior to Pavlovian training. After conditioning, liking increased for CS+_1_ and CS+_2_ relative to CS–, with ratings rising from pre- to post-learning for the CSs+ and declining for CS–, consistent with successful acquisition of the cue–outcome contingencies.

**Fig. 5.**
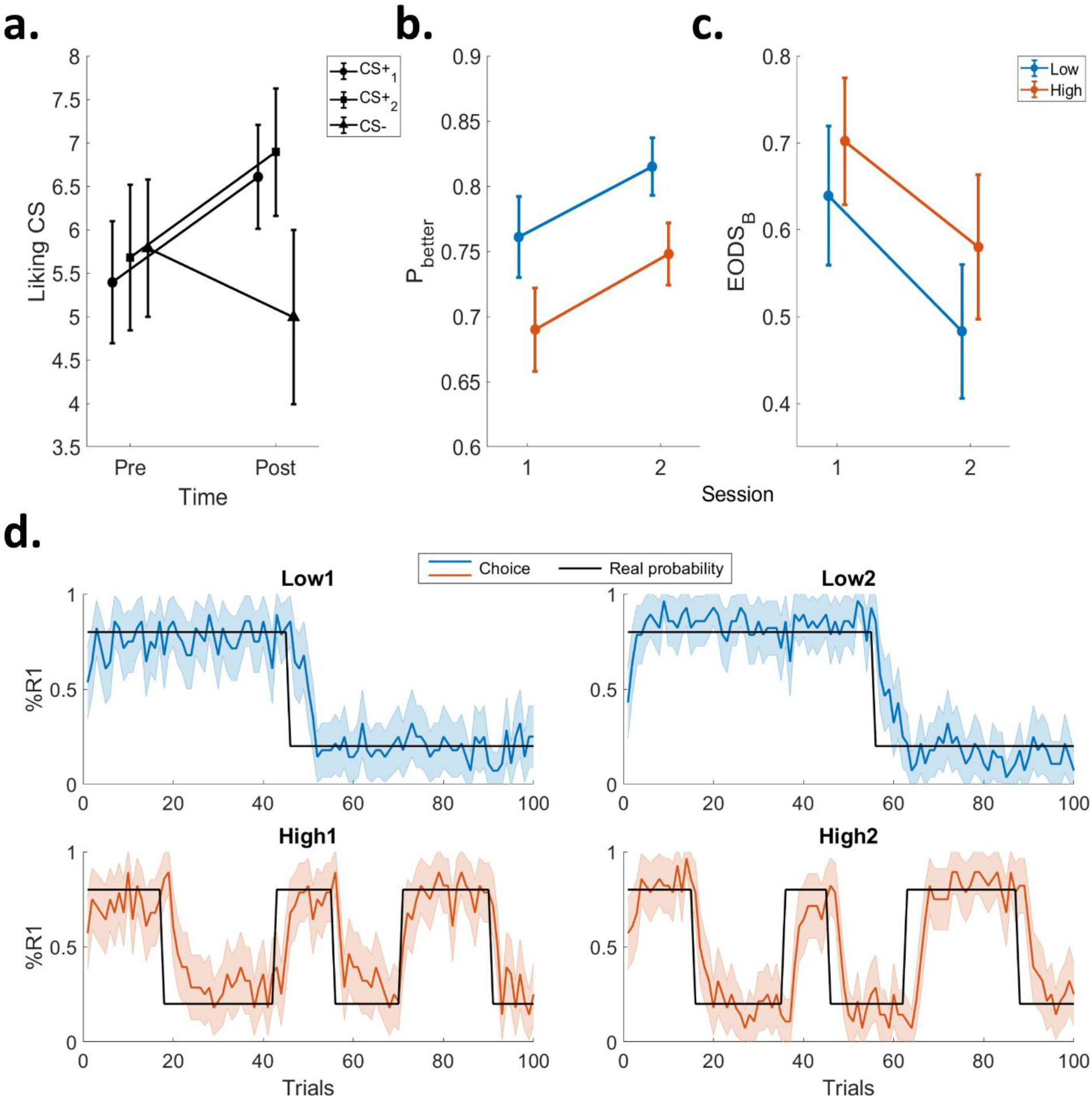
Pavlovian learning and instrumental performance. **(a)** Pavlovian conditioning. CS-liking ratings before and after conditioning for reward-predictive (CS+_1_, CS+_2_) and non-reward (CS–) cues. Ratings shifted following learning, indicating successful acquisition. Bars: group means ± 95% CI. **(b)** Instrumental accuracy. Choice accuracy (*P*_better_) by Session and Volatility. Performance improved over sessions and was higher under Low volatility. Bars: group means ± 95% CI. **(c)** Choice uncertainty (EODS*_B_*). Conditional entropy of the current choice given a better-choice response on the preceding trial. Lower entropy under Low volatility reflects more consistent selection of the optimal action. Bars: group means ± 95% CI. **(d)** Trial-by-trial choice behaviour. Percentage of R1 selections across trials. Blue and orange traces represent Low and High volatility, respectively; shaded bands indicate ± SEM. The black line denotes the true reward probability of R1.

#### Effects of environmental volatility on instrumental choice behaviour

In the instrumental learning phase of Experiment 2, all participants met the learning criterion after a minimum of two sessions. Instrumental performance, indexed by *P*better, was evaluated with a 2 × 2 repeated-measures ANOVA with Session (1 vs. 2) and Volatility (Low vs. High) as within-subject factors. The analysis revealed a main effect of Session (F_1,27_ = 18.087; p < 0.001; η_p_² = 0.401; BF₁₀ = 108.951), reflecting improved accuracy in Session 2 and a main effect of Volatility (F_1,27_ = 63.844; p < 0.001; η_p_² = 0.703; BF₁₀ = 6.344×10^4^), with superior performance in the low relative to the high volatility context (**Fig. 5b**). The Session × Volatility interaction was not significant (F_1,27_ = 0.023; p = 0.881; η_p_² = 8.507×10^-4^; BF₁₀ = 0.273). These results replicate the pattern observed in Experiment 1, indicating reduced instrumental accuracy under high volatility. A parallel 2 × 2 repeated-measures ANOVA on EODS*_B_* produced an analogous pattern: a main effect of Session (F_1,27_ = 35.812; p < 0.001; η_p_² = 0.570; BF₁₀ = 1.197×10^3^) and a main effect of Volatility (F_1,27_ = 6.274; p = 0.019; η_p_² = 0.189; BF₁₀ = 2.810), with no significant Session × Volatility interaction (F_1,27_ = 0.426; p = 0.519; η_p_² = 0.016; BF₁₀ = 0.352) (**Fig. 5c**). In line with the accuracy results, EODS*_B_* was lower in the low volatility context and higher in the high volatility context, indicating greater choice uncertainty under unstable conditions.

Trial-by-trial traces further corroborated these effects: participants performed above chance across conditions and showed increased selection of the optimal action from Session 1 to Session 2 (**Fig. 5d**), confirming a successful replication of the instrumental learning results reported for Experiment 1.

#### Computational modelling confirms adaptive learning rate modulation in volatile environments

We applied the same computational modelling framework to Experiment 2. Model comparison again identified the Rescorla-Wagner model with volatility-dependent parameters as the best-fitting model (mean BIC = 346.29; see Supplementary Table S2 for BIC values and estimated parameter distributions of all models) (**Fig. 6a**). Model validation confirmed that simulated data accurately reproduced both overall performance patterns (**Fig. 6b**) and reversal adjustment dynamics (**Fig. 6c**).

**Fig. 6.**
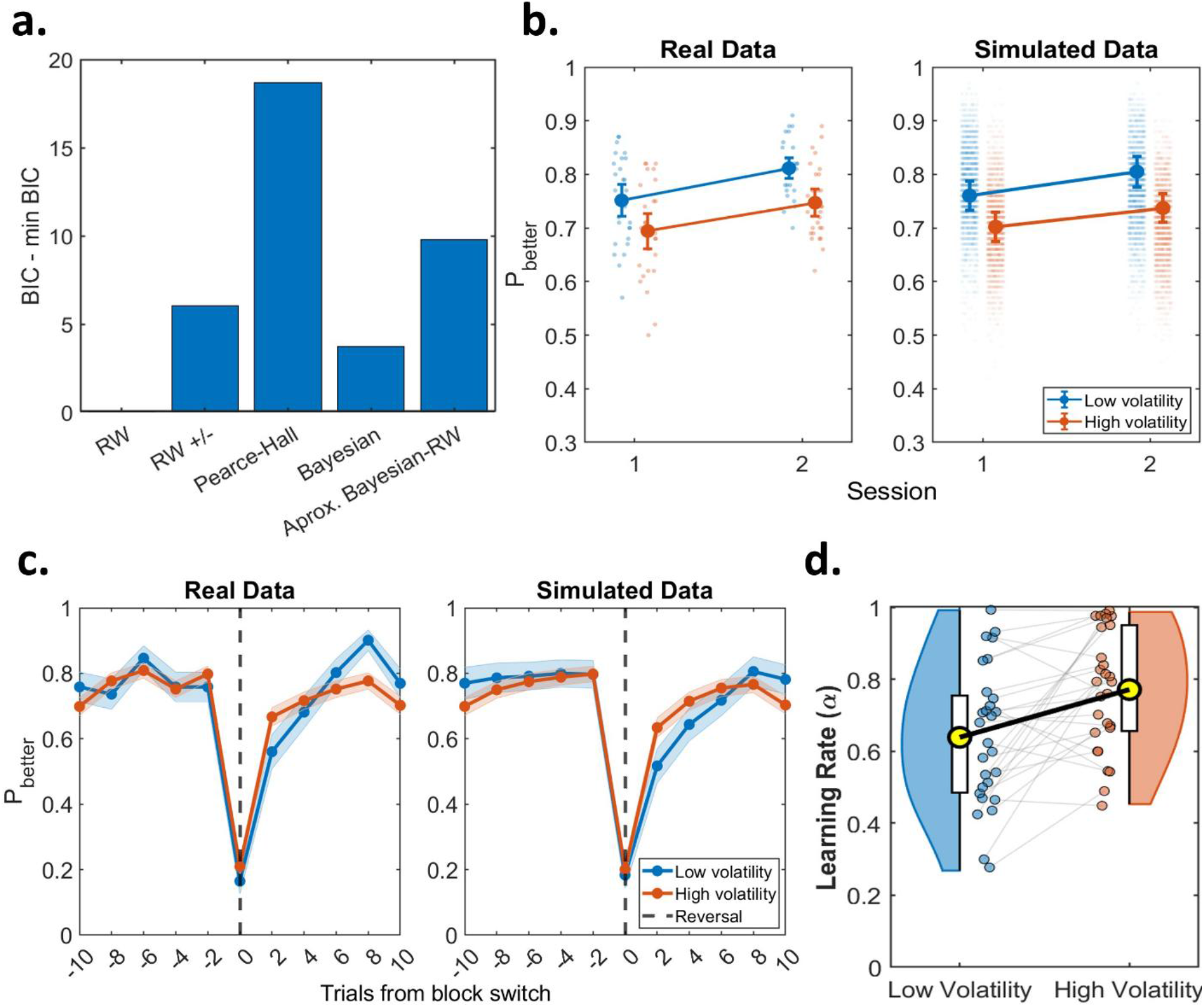
Goodness-of-fit for various computational models, model validation and model estimated parameters. **(a)** Model comparison (BIC) replicates the superiority of the Rescorla-Wagner model with volatility-dependent parameters. **(b)** Model validation: real (left) vs. simulated (right) choice accuracy across Session × Volatility. Dots = individual subjects; larger symbols with error bars = group means ± 95% CI. **(c)** Model validation: real (left) vs. simulated (right) adjustment to reversals (±10 trial window). Blue = low volatility; orange = high volatility. Colored dots with shaded regions = means ± 95% CI; vertical dotted line = reversal. **(d)** Replication of learning rate modulation: higher learning rates in high vs. low volatility contexts.

To test whether the adaptive modulation of learning rates observed in Experiment 1 are replicated in this independent sample, we performed a paired-samples *t*-test comparing learning rate parameters between low and high volatility conditions. Consistent with Experiment 1, the results revealed a significant difference in learning rates as a function of environmental volatility (*t*_27_ = – 3.481; p = 0.002; d = – 0.658; BF₁₀ = 20.993). Visual inspection of the results (**Fig. 6d**) confirmed that learning rates were higher in high volatility compared to low volatility contexts, demonstrating robust replication of the adaptive learning rate modulation across independent samples. This converging evidence further supports the conclusion that participants dynamically adjust their rate of information integration in response to environmental uncertainty. As in Experiment 1, a parallel paired-samples *t*-test on the inverse temperature parameters (β_Low_, β_High_) revealed no difference between volatility contexts (*t*₂₇ = – 0.216; *p* = 0.831; *d* = – 0.041; BF₁₀ = 0.205), confirming that the adaptation was again localized to the learning rate rather than to choice stochasticity (Supplementary Fig. S2b). Parameter recovery for the winning model was likewise satisfactory in this independent sample (all *r* > 0.81; Supplementary Fig. S3b), confirming the reliability of the parameter estimates.

To assess whether this pattern generalized to the independent sample of Experiment 2, we performed the same comparison for all five candidate models (Supplementary Tables S5, S6). As in Experiment 1, with the exception of the approximate Bayesian delta-rule model, all models showed a reliable difference in their update-rule parameter(s) between volatility contexts, consistent with the pattern observed for the winning Rescorla–Wagner model in this sample; as in Experiment 1, this reliable difference was again specific to the negative-outcome learning rate (α⁻) within the model with asymmetric learning rates, with no difference in the positive-outcome learning rate (α⁺). Inverse-temperature parameters did not differ between contexts in any model, mirroring the results reported above for the winning model (Supplementary Fig. S2b).

#### Volatility effects on Transfer without outcome feedback

To test our preregistered prediction that prior volatility alone modulates the balance of control toward the Pavlovian system, we analysed Probe trials, run without outcome feedback. For *P*better we ran a 3 (CS: CS_b_, CS_w_, CS–) × 2 (Volatility: Low, High) repeated-measures ANOVA (**Fig. 7a**). The analysis revealed a main effect of CS (F_2,54_ = 96.666; p <0.001; η_p_² = 0.782; BF₁₀ = 5.861×10^14^), a main effect of Volatility (F_1,27_ = 24.083; p < 0.001; η_p_² = 0.471; BF₁₀ = 105.800) and a significant CS × Volatility interaction (F_2,54_ = 37.366; p < 0.001; η_p_² = 0.581; BF₁₀ = 4.203×10^9^). **Fig. 7a** displays *P*better with 95% confidence intervals and offers an intuitive overview of transfer performance. First, the congruent cue (CS_b_) exerts a facilitatory effect on performance relative to the neutral control (CS–) in both volatility contexts: the CS_b_–CS− difference was reliably positive under High volatility (Δ_mean_ = 0.122, 95% CI [0.063, 0.185]) and positive under Low volatility as well, although the 95% CI closely approached zero (Δ_mean_ = 0.060, 95% CI [−0.003, 0.122]) (Supplementary Fig. S5a). Second, the neutral cue (CS–) shows no appreciable volatility-dependent shift, with largely overlapping 95% confidence intervals across contexts. Most critically, the conflict cue (CS_w_), which predicts the currently worse option, displays a volatility-dependent pattern: the CS_w_–CS− difference was reliably negative in both contexts, more than twice as large under High volatility (Δ_mean_ = −0.443, 95% CI [−0.506, −0.375]) than under Low volatility (Δ_mean_ = −0.149, 95% CI [−0.193, −0.104]) (Supplementary Fig. S5a), quantifying the strong amplification of this effect in the high-volatility context, where participants substantially shifted away from the optimal action and instead followed the cue-predicted (worse) response.

**Fig. 7.**
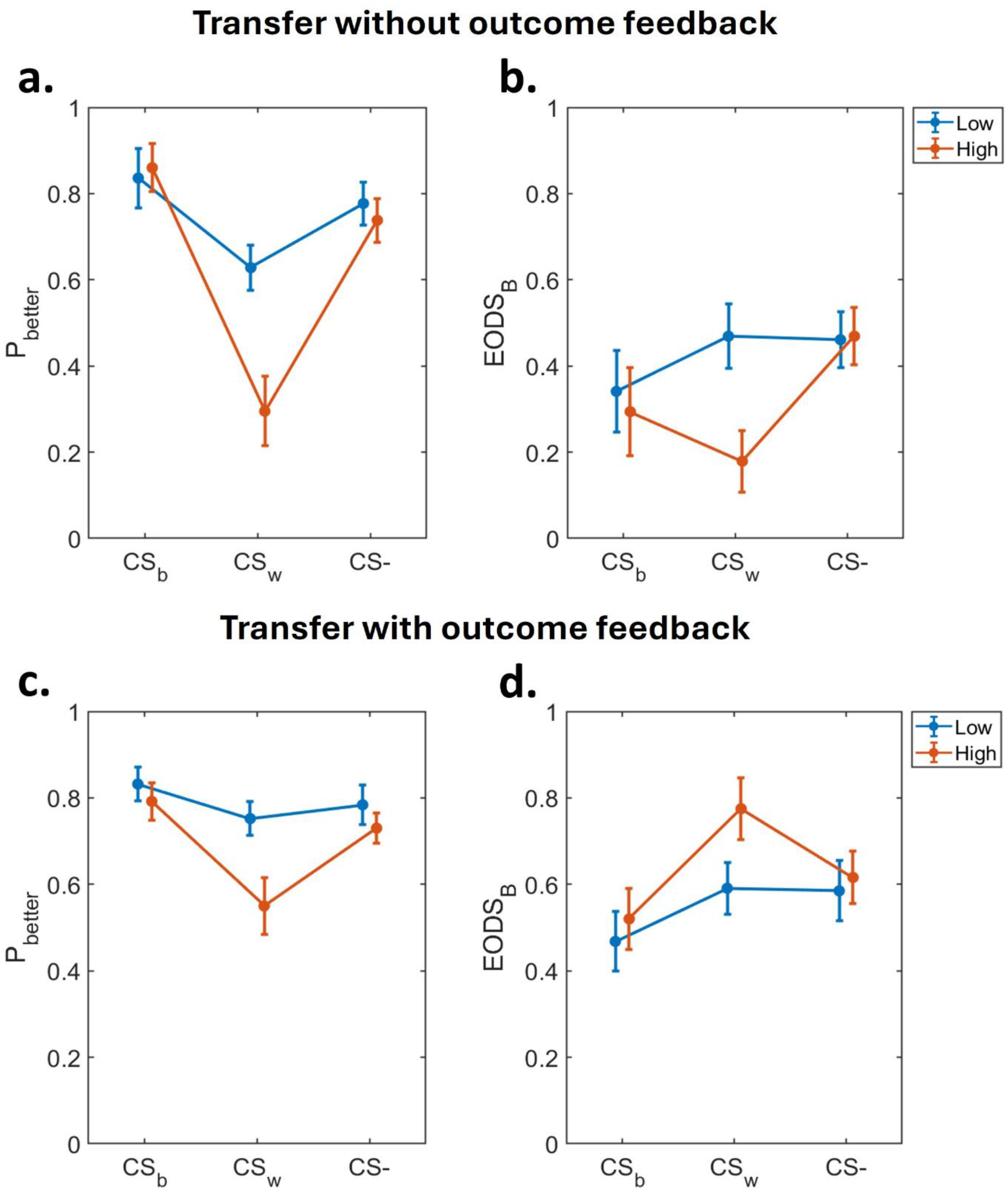
Pavlovian modulation of instrumental performance during Transfer in Experiment 2, without and with outcome feedback. **(a)** Choice accuracy (*P*_better_) without outcome feedback, by cue type and volatility context. CSb facilitates accuracy relative to CS− in both contexts; CS_w_ selectively reduces it, more strongly under high volatility. **(b)** Choice uncertainty (EODS_B_) for the same conditions. CS_b_ and CS− show stable, low entropy; CS_w_ entropy decreases under high volatility, indicating a highly consistent, cue-driven choice policy. **(c)** Choice accuracy (*P*_better_) with outcome feedback, by cue type and volatility context. The pattern replicates Experiment 1: CS_b_ facilitates accuracy; CS_w_ selectively reduces it under high volatility. **(d)** Choice uncertainty (EODS_B_) for the same conditions. CS_b_ and CS−remain stable; CS_w_ entropy increases under high volatility, indicating continued competition between the two systems. Error bars represent 95% confidence intervals.

As in the previous experiment, we complemented the accuracy analysis with a 3 (CS: CS_b_, CS_w_, CS–) × 2 (Volatility: Low, High) repeated-measures ANOVA on EODS*_B_* (**Fig. 7b**). This analysis revealed a significant main effect of Volatility (F_1,27_ = 16.191; p < 0.001; η_p_² = 0.375; BF₁₀ = 11.183), CS (F_2,54_ = 11.127; p < 0.001; η_p_² = 0.292; BF₁₀ = 57.689) and a significant Volatility × CS interaction (F_2,54_= 9.507; p < 0.001; η_p_² = 0.260; BF₁₀ = 2.168×10^3^).

Specifically, a visual inspection of the results shows that CS_b_ consistently elicits low entropy regardless of volatility, indicating stable better-choice behaviour when the cue matches the currently better action. The CS– likewise shows overlapping confidence intervals across volatility, suggesting no systematic volatility-dependent change for the control condition. By contrast, the CS_w_ displays a pronounced volatility-dependent shift, showing a pattern opposite to that observed in Experiment 1. Indeed, Under Low volatility, choice entropy on conflict trials is comparable to the neutral control, indicating no reliable cue-induced modulation of choice uncertainty. Critically, under high volatility, entropy is markedly reduced relative both to the neutral condition (CS–) and to the same condition (CS_w_) in the low-volatility context. This reduction reflects that participants adopt a highly consistent, rather than uncertain, choice policy; however, this consistency is systematically aligned with the Pavlovian-predicted action, which is currently worse option.

In summary, performance remains unaffected by contextual volatility in the presence of CS_b_ and CS– conditions, whereas prior exposure to a volatile environment selectively amplifies Pavlovian bias when cues predict the currently worse option (CS_w_). Moreover, this effect is corroborated by a reduction in choice entropy on CS_w_ trials, indicating a highly consistent (yet suboptimal) choice policy when Pavlovian cues predict the currently worse option. Crucially, this pattern emerges without outcome feedback, indicating that Pavlovian control over choice is shaped by learned environmental volatility rather than sustained by continuous reinforcement.

#### Volatility effects on Transfer with outcome feedback

To assess transfer performance when outcome information was available, we analysed choices during the Normal segments of the Transfer phase (feedback and point counter visible). A 3 × 2 repeated-measures ANOVA on *P*better revealed a significant main effects of CS (F_2,54_ = 31.345; p <0.001; η_p_² = 0.537; BF₁₀ = 2.869×10^6^), Volatility (F_1,27_ = 27.676; p <0.001; η_p_² = 0.506; BF₁₀ = 601.279) and a significant CS × Volatility interaction (F_2,54_ = 11.970; p <0.001; η_p_² = 0.307; BF₁₀ = 4.651×10^3^). Visual inspection of the results (**Fig. 7c**) indicates a pattern that closely replicates the Transfer phase results observed in Experiment 1. Across volatility contexts, performance is consistently higher following low- than high-volatility contexts; CS_b_ facilitates optimal responding relative to the neutral control cue (CS–): the CS_b_–CS− difference was reliably positive under High volatility (Δ_mean_ = 0.060, 95% CI [0.004, 0.117]) and positive under Low volatility as well, with a 95% CI that only narrowly excluded zero (Δ_mean_ = 0.048, 95% CI [0.001, 0.097]). In contrast, the impact of the conflict cue (CS_w_) depended strongly on environmental volatility: the CS_w_–CS− difference was centred near zero under Low volatility (Δ_mean_ = −0.032, 95% CI [−0.072, 0.008]), but reliably negative under High volatility (Δ_mean_ = −0.182, 95% CI [−0.236, −0.125]) (Supplementary Fig. S5b).

These findings were further corroborated by the analysis of EODS*_B_*. A repeated-measures ANOVA revealed significant main effects of Volatility (F_1,27_ = 9.394; p = 0.005; η_p_² = 0.258; BF₁₀ = 7.088) and CS (F_2,54_ = 14.157; p < 0.001; η_p_² = 0.344; BF₁₀ = 1.395×10^3^) and a significant Volatility x CS interaction (F_2,54_ = 3.406; p = 0.040; η_p_²= 0.112; BF₁₀ = 2.250). Critically, the interaction was driven specifically by the CS_w_. While the influence of both the CS_b_ and the neutral control CS– remained stable across volatility conditions, a difference emerged for CS_w_ (**Fig. 7d**). In the high volatility context, CS_w_ induced significantly greater uncertainty regarding the better choice compared to low volatility, indicating that environmental volatility enhances Pavlovian interference on choice policy even when outcome feedback is explicitly provided.

In the Normal segments, where outcome feedback was available, the same pattern observed in Experiment 1 emerged: CS_b_ facilitated performance relative to CS− across volatility contexts, while CS_w_ selectively reduced *P*better and increased EODS*_B_* under high volatility, producing a significant CS × Volatility interaction for both measures.

## Discussion

Across two preregistered experiments, we investigated how environmental volatility shapes the balance between instrumental and Pavlovian control over choice. In both experiments, participants first performed an instrumental learning task in which two actions led to different outcomes, and where the rate of contingency reversals defined a low- and a high-volatility context. They were then tested in a Transfer phase, choosing between the same two actions in the presence of reward-predictive cues. The two experiments differed in whether outcome feedback was available during Transfer. In Experiment 1, choices were made with outcome feedback throughout, so both systems could continue to guide behaviour. In Experiment 2, the Transfer phase alternated segments with outcome feedback and segments without outcome feedback, the latter allowing us to measure Pavlovian influence while participants could still track the current volatility context. This lets us test whether the influence of volatile learning depends on ongoing reinforcement or persists without outcome feedback.

Three main findings emerged. First, participants adjusted to volatility as expected, increasing their learning rate in the high-volatility context. Second, during Transfer, high volatility increased cue-driven choice, but only when the cue and the instrumental system favoured different actions. Third, the entropy of choice strategy moved in opposite directions across experiments, rising under conflict when feedback was available and falling when it was not. Together, these results indicate that volatility shifts the balance of control toward Pavlovian cues and that this shift affects behaviour only when the two systems conflict.

### Volatility shapes instrumental learning

To establish that volatility altered learning itself, we fitted computational models to the instrumental learning phase alone, before any Pavlovian cues were introduced, testing whether volatility changed participants’ learning dynamics, independently of how cues would later influence choice. Participants learned with a higher learning rate, weighting recent outcomes more heavily, in the high- than in the low-volatility context and this effect was specific across both experiments. Crucially, choice stochasticity did not differ between contexts, so what changed was the speed of expectation updating, not choice noise. This matches what is consistently reported for learning under volatility (Behrens et al., 2007; Massi et al., 2018; Nassar et al., 2010; Payzan-LeNestour & Bossaerts, 2011).

However, faster updating came at a cost: accuracy was lower and choice entropy higher under high volatility. When the optimal action keeps changing, even accurate learning cannot yield consistently correct choice and the instrumental system remains persistently uncertain about which action is currently better, a defining feature of unexpected uncertainty (Bland & Schaefer, 2012; Soltani & Izquierdo, 2019; Soltani & Koechlin, 2022; Yu & Dayan, 2005).

### With outcome feedback, the two systems compete

This loss of instrumental informativeness set the stage for the Transfer phase, where we measured how Pavlovian cues shaped choice. When a cue predicted the outcome of the currently worse action (CS_w_), participants in the high-volatility context chose that action more often and accuracy dropped. When the cue predicted the currently better action (CS_b_), performance was preserved: CS_b_ yielded the highest accuracy of any condition, in both the low- and high-volatility context. The same asymmetry appeared across both experiments whenever choices were made with outcome feedback.

These two effects are the opposite faces of a single process. On each trial, the action predicted by the cue is weighted against the action favoured by instrumental learning. When the two agree (CS_b_), the converging predictions reinforce one another and choice accuracy increases. When they disagree (CS_w_), the cue weighs more heavily as instrumental precision declines under volatility and choice accuracy decreases. CS_b_ and CS_w_, therefore, arise from the same mechanism, differing only in whether the cue and the instrumental estimate happen to converge or conflict. This is why neither volatility nor conflict was sufficient on its own. Volatility alone produced no cost when the two systems agreed: under CS_b_ and CS−, participants in the high-volatility context performed as well as those in the low-volatility one. And competition between the two systems was equally insufficient without volatility: under low volatility, CS_w_ did not depart from neutral control (CS−), because instrumental knowledge stayed informative enough to override the cue. Only when both held together (i) a degraded instrumental system and (ii) a cue placing the two systems in conflict did the balance of control shift toward cue-driven, suboptimal choice.

The entropy of choice strategy (EODS*_B_*) confirms what kind of change this was. With outcome feedback available, conflict increased entropy: under CS_w_ in the high-volatility context, choice became more variable. Participants alternated between the instrumental and Pavlovian systems rather than committing to either and this variability is the result of two systems competing for control (Dorfman & Gershman, 2019; Guitart-Masip et al., 2012; Huys et al., 2012).

Our findings extend what is known about when Pavlovian control takes over. Prior work has shown that it grows when an environment is uncontrollable, when outcomes stop following from actions at all (Dorfman & Gershman, 2019; Gershman et al., 2021; Huys & Dayan, 2009). However, controllability and volatility are separate things and recent work has shown they can be estimated independently (Piray & Daw, 2020, 2021): an environment can be fully controllable and still highly volatile. That is the case in our task. Actions always led to their outcomes, and participants kept control over what happened throughout. What changed was something narrower: which action was the better one and how often that changed.

### Without outcome feedback, the cue captures control

Removing outcome feedback changed the nature of the effect. During the segments without outcome feedback, accuracy under CS_w_ again fell after high-volatility learning, showing that the effect does not depend on ongoing reinforcement. Crucially, this time entropy fell alongside it: choice became highly consistent while accuracy declined. Without outcome feedback to update on, the instrumental system had nothing to contribute and the cue was left as the sole source of value; so choice shifted onto a single, cue-driven policy (Cartoni et al., 2013; Dayan et al., 2006; Dayan & Berridge, 2014).

Read against Experiment 1, this confirms the dissociation. The same conflict that made choice more variable when both systems were active made it uniform when only the cue remained. This dissociation in entropy points to a qualitative change in which system controls choice, from two systems competing to one taking command (Woo et al., 2025b). It is difficult to reconcile with the idea that volatility simply adds noise, which would increase entropy in both experiments rather than move it in opposite directions.

The effect held even without outcome feedback, indicating that it does not depend on ongoing reinforcement. Thus, what participants carried into Transfer was the context-linked learning-rate signature established during instrumental learning, plausibly reinstated by the contextual cues (background colour) that had marked each volatility context (Bouton, 2004; Gershman et al., 2010; Radulescu et al., 2021).

### Limitations and future directions

Our data establish that learned volatility shifts the balance of control toward Pavlovian responding, but because our computational models captured only the instrumental system, they cannot identify the mechanism through which this shift occurs. At least two possibilities remain compatible with the present results: volatility may reduce the weight assigned to the instrumental controller, increasing the relative influence of an unchanged Pavlovian signal (Dorfman & Gershman, 2019; Gershman et al., 2021; S. W. Lee et al., 2014), or it may enhance the rate of Pavlovian value updating itself (Degni et al., 2026). Both could be happening together and adjudicating between them will require models that represent both controllers (i.e. Pavlovian and instrumental) concurrently across all task phases, a step beyond the scope of the present study.

These findings carry implications for cue-driven behavior in contexts marked by uncertainty. Exaggerated Pavlovian control over choice has been implicated in addiction, gambling and compulsive disorders (Everitt & Robbins, 2005; Marzuki et al., 2024; Sommer et al., 2017), conditions frequently associated with volatile, unpredictable circumstances. Our results suggest that environmental volatility may itself contribute to this shift toward cue-driven responding, independently of any change in controllability, a mechanism worth examining alongside other established vulnerability factors in these conditions.

### Conclusions

Together, these findings show that environmental volatility shifts the balance of control toward Pavlovian cues through a precise, condition-dependent arbitration mechanism, and that the entropy of choice strategy distinguishes competition between systems from capture by a single system.

## Bibliography

Andraszewicz, S., Scheibehenne, B., Rieskamp, J., Grasman, R., Verhagen, J., & Wagenmakers, E.-J. (2015). An Introduction to Bayesian Hypothesis Testing for Management Research. Journal of Management, 41(2), 521–543. 10.1177/0149206314560412

Badioli, M., Degni, L. A. E., Dalbagno, D., Danti, C., Starita, F., Di Pellegrino, G., Benassi, M., & Garofalo, S. (2024). Unraveling the influence of Pavlovian cues on decision-making: A pre-registered meta-analysis on Pavlovian-to-instrumental transfer. Neuroscience & Biobehavioral Reviews, 164, 105829. 10.1016/j.neubiorev.2024.105829

Behrens, T. E. J., Woolrich, M. W., Walton, M. E., & Rushworth, M. F. S. (2007). Learning the value of information in an uncertain world. Nature Neuroscience, 10(9), 1214–1221. 10.1038/nn1954

Blain, B., & Rutledge, R. B. (2020). Momentary subjective well-being depends on learning and not reward. eLife, 9, e57977. 10.7554/eLife.57977

Bland, A. R., & Schaefer, A. (2012). Different Varieties of Uncertainty in Human Decision-Making. Frontiers in Neuroscience, 6. 10.3389/fnins.2012.00085

Boureau, Y.-L., Sokol-Hessner, P., & Daw, N. D. (2015). Deciding How To Decide: Self-Control and Meta-Decision Making. Trends in Cognitive Sciences, 19(11), 700–710. 10.1016/j.tics.2015.08.013

Bouton, M. E. (2004). Context and Behavioral Processes in Extinction: Table 1. Learning & Memory, 11(5), 485–494. 10.1101/lm.78804

Calin-Jageman, R. J., & Cumming, G. (2019). The New Statistics for Better Science: Ask How Much, How Uncertain, and What Else Is Known. The American Statistician, 73(sup1), 271–280. 10.1080/00031305.2018.1518266

Campbell, J. I. D., & Thompson, V. A. (2012). MorePower 6.0 for ANOVA with relational confidence intervals and Bayesian analysis. Behavior Research Methods, 44(4), 1255–1265. 10.3758/s13428-012-0186-0

Cartoni, E., Moretta, T., Puglisi-Allegra, S., Cabib, S., & Baldassarre, G. (2015). The Relationship Between Specific Pavlovian Instrumental Transfer and Instrumental Reward Probability. Frontiers in Psychology, 6. 10.3389/fpsyg.2015.01697

Cartoni, E., Puglisi-Allegra, S., & Baldassarre, G. (2013). The three principles of action: A Pavlovian-instrumental transfer hypothesis. Frontiers in Behavioral Neuroscience, 7. 10.3389/fnbeh.2013.00153

Courville, A. C., Daw, N. D., & Touretzky, D. S. (2006). Bayesian theories of conditioning in a changing world. Trends in Cognitive Sciences, 10(7), 294–300. 10.1016/j.tics.2006.05.004

Cumming, G. (2014). The New Statistics: Why and How. Psychological Science, 25(1), 7–29. 10.1177/0956797613504966

Daw, N. D., Kakade, S., & Dayan, P. (2002). Opponent interactions between serotonin and dopamine. Neural Networks, 15(4–6), 603–616. 10.1016/S0893-6080(02)00052-7

Daw, N. D., Niv, Y., & Dayan, P. (2005). Uncertainty-based competition between prefrontal and dorsolateral striatal systems for behavioral control. Nature Neuroscience, 8(12), 1704–1711. 10.1038/nn1560

Dayan, P., & Berridge, K. C. (2014). Model-based and model-free Pavlovian reward learning: Revaluation, revision, and revelation. Cognitive, Affective, & Behavioral Neuroscience, 14(2), 473–492. 10.3758/s13415-014-0277-8

Dayan, P., Niv, Y., Seymour, B., & D. Daw, N. (2006). The misbehavior of value and the discipline of the will. Neural Networks, 19(8), 1153–1160. 10.1016/j.neunet.2006.03.002

Degni, L. A. E., Dalbagno, D., Starita, F., Benassi, M., Di Pellegrino, G., & Garofalo, S. (2022). General Pavlovian-to-instrumental transfer in humans: Evidence from Bayesian inference. Frontiers in Behavioral Neuroscience, 16, 945503. 10.3389/fnbeh.2022.945503

Degni, L. A. E., Danti, C., Finotti, G., Starita, F., Di Pellegrino, G., & Garofalo, S. (2025). Pavlovian bias instigates suboptimal choices in humans. Behaviour Research and Therapy, 195, 104906. 10.1016/j.brat.2025.104906

Degni, L. A. E., Garofalo, S., Finotti, G., Starita, F., Robbins, T. W., & Di Pellegrino, G. (2024). Sex differences in motivational biases over instrumental actions. Npj Science of Learning, 9(1), 62. 10.1038/s41539-024-00246-6

Degni, L. A. E., Mattioni, L., Danti, C., Bernardi, V., Finotti, G., Badioli, M., Starita, F., Soltani, A., Di Pellegrino, G., & Garofalo, S. (2026). Reduced Pavlovian Value Updating Alters Decision-Making in Sign-Trackers. The Journal of Neuroscience, 46(3), e1465252025. 10.1523/JNEUROSCI.1465-25.2025

Dorfman, H. M., & Gershman, S. J. (2019). Controllability governs the balance between Pavlovian and instrumental action selection. Nature Communications, 10(1), 5826. 10.1038/s41467-019-13737-7

Everitt, B. J., & Robbins, T. W. (2005). Neural systems of reinforcement for drug addiction: From actions to habits to compulsion. Nature Neuroscience, 8(11), 1481–1489. 10.1038/nn1579

Farashahi, S., Donahue, C. H., Hayden, B. Y., Lee, D., & Soltani, A. (2019). Flexible combination of reward information across primates. Nature Human Behaviour, 3(11), 1215–1224. 10.1038/s41562-019-0714-3

Farashahi, S., Rowe, K., Aslami, Z., Lee, D., & Soltani, A. (2017). Feature-based learning improves adaptability without compromising precision. Nature Communications, 8(1), 1768. 10.1038/s41467-017-01874-w

Frank, M. J., Seeberger, L. C., & O’Reilly, R. C. (2004). By Carrot or by Stick: Cognitive Reinforcement Learning in Parkinsonism. Science, 306(5703), 1940–1943. 10.1126/science.1102941

Garofalo, S., Battaglia, S., Starita, F., & di Pellegrino, G. (2021). Modulation of cue-guided choices by transcranial direct current stimulation. Cortex, 137, 124–137. 10.1016/j.cortex.2021.01.004

Garofalo, S., & di Pellegrino, G. (2015). Individual differences in the influence of task-irrelevant Pavlovian cues on human behavior. Frontiers in Behavioral Neuroscience, 9. 10.3389/fnbeh.2015.00163

Gershman, S. J. (2015). A Unifying Probabilistic View of Associative Learning. PLOS Computational Biology, 11(11), e1004567. 10.1371/journal.pcbi.1004567

Gershman, S. J. (2016). Empirical priors for reinforcement learning models. Journal of Mathematical Psychology, 71, 1–6. 10.1016/j.jmp.2016.01.006

Gershman, S. J., Blei, D. M., & Niv, Y. (2010). Context, learning, and extinction. Psychological Review, 117(1), 197–209. 10.1037/a0017808

Gershman, S. J., Guitart-Masip, M., & Cavanagh, J. F. (2021). Neural signatures of arbitration between Pavlovian and instrumental action selection. PLOS Computational Biology, 17(2), e1008553. 10.1371/journal.pcbi.1008553

Guitart-Masip, M., Huys, Q. J. M., Fuentemilla, L., Dayan, P., Duzel, E., & Dolan, R. J. (2012). Go and no-go learning in reward and punishment: Interactions between affect and effect. NeuroImage, 62(1), 154–166. 10.1016/j.neuroimage.2012.04.024

Ho, J., Tumkaya, T., Aryal, S., Choi, H., & Claridge-Chang, A. (2019). Moving beyond P values: Data analysis with estimation graphics. Nature Methods, 16(7), 565–566. 10.1038/s41592-019-0470-3

Hopkins, K. D., & Weeks, D. L. (1990). Tests for Normality and Measures of Skewness and Kurtosis: Their Place in Research Reporting. Educational and Psychological Measurement, 50(4), 717–729. 10.1177/0013164490504001

Huys, Q. J. M., Cools, R., Gölzer, M., Friedel, E., Heinz, A., Dolan, R. J., & Dayan, P. (2011). Disentangling the Roles of Approach, Activation and Valence in Instrumental and Pavlovian Responding. PLoS Computational Biology, 7(4), e1002028. 10.1371/journal.pcbi.1002028

Huys, Q. J. M., & Dayan, P. (2009). A Bayesian formulation of behavioral control. Cognition, 113(3), 314–328. 10.1016/j.cognition.2009.01.008

Huys, Q. J. M., Eshel, N., O’Nions, E., Sheridan, L., Dayan, P., & Roiser, J. P. (2012). Bonsai Trees in Your Head: How the Pavlovian System Sculpts Goal-Directed Choices by Pruning Decision Trees. PLoS Computational Biology, 8(3), e1002410. 10.1371/journal.pcbi.1002410

Kruschke, J. K. (2021). Bayesian Analysis Reporting Guidelines. Nature Human Behaviour, 5(10), 1282–1291. 10.1038/s41562-021-01177-7

Lakens, D. (2013). Calculating and reporting effect sizes to facilitate cumulative science: A practical primer for t-tests and ANOVAs. Frontiers in Psychology, 4. 10.3389/fpsyg.2013.00863

Lee, M. D., & Wagenmakers, E.-J. (2014). Bayesian Cognitive Modeling: A Practical Course (1^a^ ed.). Cambridge University Press. 10.1017/CBO9781139087759

Lee, S. W., Shimojo, S., & O’Doherty, J. P. (2014). Neural Computations Underlying Arbitration between Model-Based and Model-free Learning. Neuron, 81(3), 687–699. 10.1016/j.neuron.2013.11.028

Love, J., Selker, R., Marsman, M., Jamil, T., Dropmann, D., Verhagen, J., Ly, A., Gronau, Q. F., Smíra, M., Epskamp, S., Matzke, D., Wild, A., Knight, P., Rouder, J. N., Morey, R. D., & Wagenmakers, E.-J. (2019). **JASP**: Graphical Statistical Software for Common Statistical Designs. Journal of Statistical Software, 88(2). 10.18637/jss.v088.i02

Marzuki, A. A., Banca, P., Garofalo, S., Degni, L. A. E., Dalbagno, D., Badioli, M., Sule, A., Kaser, M., Conway-Morris, A., Sahakian, B. J., & Robbins, T. W. (2024). Compulsive avoidance in youths and adults with OCD: An aversive pavlovian-to-instrumental transfer study. Translational Psychiatry, 14(1), 308. 10.1038/s41398-024-03028-1

Massi, B., Donahue, C. H., & Lee, D. (2018). Volatility Facilitates Value Updating in the Prefrontal Cortex. Neuron, 99(3), 598–608.e4. 10.1016/j.neuron.2018.06.033

Mathôt, S., Schreij, D., & Theeuwes, J. (2012). OpenSesame: An open-source, graphical experiment builder for the social sciences. Behavior Research Methods, 44(2), 314–324. 10.3758/s13428-011-0168-7

Mathys, C. D., Lomakina, E. I., Daunizeau, J., Iglesias, S., Brodersen, K. H., Friston, K. J., & Stephan, K. E. (2014). Uncertainty in perception and the Hierarchical Gaussian Filter. Frontiers in Human Neuroscience, 8. 10.3389/fnhum.2014.00825

McGuire, J. T., Nassar, M. R., Gold, J. I., & Kable, J. W. (2014). Functionally Dissociable Influences on Learning Rate in a Dynamic Environment. Neuron, 84(4), 870–881. 10.1016/j.neuron.2014.10.013

Nassar, M. R., Rumsey, K. M., Wilson, R. C., Parikh, K., Heasly, B., & Gold, J. I. (2012). Rational regulation of learning dynamics by pupil-linked arousal systems. Nature Neuroscience, 15(7), 1040–1046. 10.1038/nn.3130

Nassar, M. R., Wilson, R. C., Heasly, B., & Gold, J. I. (2010). An Approximately Bayesian Delta-Rule Model Explains the Dynamics of Belief Updating in a Changing Environment. The Journal of Neuroscience, 30(37), 12366–12378. 10.1523/JNEUROSCI.0822-10.2010

Palminteri, S., Wyart, V., & Koechlin, E. (2017). The Importance of Falsification in Computational Cognitive Modeling. Trends in Cognitive Sciences, 21(6), 425–433. 10.1016/j.tics.2017.03.011

Payzan-LeNestour, E., & Bossaerts, P. (2011). Risk, Unexpected Uncertainty, and Estimation Uncertainty: Bayesian Learning in Unstable Settings. PLoS Computational Biology, 7(1), e1001048. 10.1371/journal.pcbi.1001048

Pearce, J. M., & Hall, G. (1980). A model for Pavlovian learning: Variations in the effectiveness of conditioned but not of unconditioned stimuli. Psychological Review, 87(6), 532–552. 10.1037/0033-295X.87.6.532

Piray, P., & Daw, N. D. (2020). A simple model for learning in volatile environments. PLOS Computational Biology, 16(7), e1007963. 10.1371/journal.pcbi.1007963

Piray, P., & Daw, N. D. (2021). A model for learning based on the joint estimation of stochasticity and volatility. Nature Communications, 12(1), 6587. 10.1038/s41467-021-26731-9

Radulescu, A., Shin, Y. S., & Niv, Y. (2021). Human Representation Learning. Annual Review of Neuroscience, 44(1), 253–273. 10.1146/annurev-neuro-092920-120559

Rangel, A., Camerer, C., & Montague, P. R. (2008). A framework for studying the neurobiology of value-based decision making. Nature Reviews Neuroscience, 9(7), 545–556. 10.1038/nrn2357

Roesch, M. R., Esber, G. R., Li, J., Daw, N. D., & Schoenbaum, G. (2012). Surprise! Neural correlates of Pearce–Hall and Rescorla–Wagner coexist within the brain. European Journal of Neuroscience, 35(7), 1190–1200. 10.1111/j.1460-9568.2011.07986.x

Shannon, C. E. (1948). A Mathematical Theory of Communication. Bell System Technical Journal, 27(3), 379–423. 10.1002/j.1538-7305.1948.tb01338.x

Soltani, A., & Izquierdo, A. (2019). Adaptive learning under expected and unexpected uncertainty. Nature Reviews Neuroscience, 20(10), 635–644. 10.1038/s41583-019-0180-y

Soltani, A., & Koechlin, E. (2022). Computational models of adaptive behavior and prefrontal cortex. Neuropsychopharmacology, 47(1), 58–71. 10.1038/s41386-021-01123-1

Soltani, A., Murray, J. D., Seo, H., & Lee, D. (2021). Timescales of cognition in the brain. Current Opinion in Behavioral Sciences, 41, 30–37. 10.1016/j.cobeha.2021.03.003

Sommer, C., Garbusow, M., Jünger, E., Pooseh, S., Bernhardt, N., Birkenstock, J., Schad, D. J., Jabs, B., Glöckler, T., Huys, Q. M., Heinz, A., Smolka, M. N., & Zimmermann, U. S. (2017). Strong seduction: Impulsivity and the impact of contextual cues on instrumental behavior in alcohol dependence. Translational Psychiatry, 7(8), e1183–e1183. 10.1038/tp.2017.158

Sutton, R. S., & Barto, A. (2018). Reinforcement learning: An introduction (Second edition). The MIT Press.

Swart, J. C., Froböse, M. I., Cook, J. L., Geurts, D. E., Frank, M. J., Cools, R., & Den Ouden, H. E. (2017). Catecholaminergic challenge uncovers distinct Pavlovian and instrumental mechanisms of motivated (in)action. eLife, 6, e22169. 10.7554/eLife.22169

Trepka, E., Spitmaan, M., Bari, B. A., Costa, V. D., Cohen, J. Y., & Soltani, A. (2021). Entropy-based metrics for predicting choice behavior based on local response to reward. Nature Communications, 12(1), 6567. 10.1038/s41467-021-26784-w

Wagner, A. R., & Rescorla, R. A. (1972). Inhibition in Pavlovian conditioning: Application of a theory. In Inhibition and learning (pp. 301–336). Academic Press Inc.

Wilson, R. C., & Collins, A. G. (2019). Ten simple rules for the computational modeling of behavioral data. eLife, 8, e49547. 10.7554/eLife.49547

Wilson, R. C., & Niv, Y. (2012). Inferring Relevance in a Changing World. Frontiers in Human Neuroscience, 5. 10.3389/fnhum.2011.00189

Woo, J. H., Aguirre, C. G., Bari, B. A., Tsutsui, K.-I., Grabenhorst, F., Cohen, J. Y., Schultz, W., Izquierdo, A., & Soltani, A. (2023). Mechanisms of adjustments to different types of uncertainty in the reward environment across mice and monkeys. *Cognitive, Affective*, & Behavioral Neuroscience, 23(3), 600–619. 10.3758/s13415-022-01059-z

Woo, J. H., Balaji, L., & Soltani, A. (2025a). An Information-Theoretic Framework for Understanding Learning and Choice Under Uncertainty. Entropy, 27(10), 1056. 10.3390/e27101056

Woo, J. H., Costa, V. D., Taswell, C. A., Rothenhoefer, K. M., Averbeck, B. B., & Soltani, A. (2025b). Contribution of amygdala to dynamic model arbitration under uncertainty. Nature Communications, 16(1), 11704. 10.1038/s41467-025-66745-1

Yu, A. J., & Dayan, P. (2005). Uncertainty, Neuromodulation, and Attention. Neuron, 46(4), 681–692. 10.1016/j.neuron.2005.04.026

